# Testing the Growth Rate and Temperature Compensation Hypotheses in Marine Bacterioplankton

**DOI:** 10.1101/2022.06.28.497896

**Authors:** Shira Givati, Xingyu Yang, Daniel Sher, Eyal Rahav

## Abstract

Two different hypotheses have been raised as to how temperature affects resource allocation in microorganisms. The translation-compensation hypothesis (TCH) predicts that the increase in enzymatic efficiency with temperature results in fewer required ribosomes per cell and lower RNA:protein ratio. In contrast, the growth rate hypothesis (GRH) predicts that increasing growth rate with temperature requires more ribosomes and hence a higher cellular RNA:protein. We tested these two hypotheses in lab cultures of *Prochlorococcus* and *Alteromonas* as well as over an annual cycle in the Eastern Mediterranean Sea. The RNA:protein of *Alteromonas* mostly decreased with temperature in accordance with the TCH, while that of *Prochlorococcus* increased with temperature, as predicted by the GRH. No support was found for either hypotheses in surface waters from the Eastern Mediterranean, whereas the fraction of phosphorus in RNA was positively correlated with per-cell bacterial production in the deep chlorophyll maximum, supporting the GRH in this niche. A considerable part of the cellular phosphorus was not allocated to RNA, DNA, phospholipids or polyphosphate, raising the question which cellular molecules contain these P reserves. While macromolecular quotas differed significantly between laboratory cultures and field samples, these were connected through a power law, suggesting common rules of resource allocation.

**Originality-Significance statement:** We investigated whether the translation-compensation hypothesis (TCH) or growth rate hypothesis (GRH) affect the macromolecular composition and phosphorus allocation in both lab cultures of *Prochlorococcus* and *Alteromonas* as well as in seawater with natural microbial communities. Our results highlight that the TCH and GRH may each be applicable to different organisms (autotroph or heterotroph), physiological states or environmental conditions. Testing the applicability of theoretical models such as the TCH and GRH in lab cultures and field samples is an important step toward mechanistic models of bacterial physiology. This is especially important to our understanding of how bacterioplankton allocate resources in response to changes in environmental conditions such as temperature and nutrient stress, which are likely to expand due to the predicted global changes.

## Introduction

Phytoplankton and heterotrophic bacteria drive the oceanic biogeochemical cycles of carbon (C), nitrogen (N) and phosphorous (P) (Litchman *et al*., 2015). Their physiology is affected by different cellular and environmental conditions such as growth rate (Sterner and Elser, 2002), the organism’s dietary needs (Martiny et al., 2016; Quigg et al., 2003), available nutrients supply (Moore et al., 2013; Reich & Oleksyn, 2004), temperature (Toseland *et al*., 2013), and biotic interactions (Ankrah *et al*., 2014; Aharonovich and Sher, 2016; Biller *et al*., 2018). These changes can then affect the resource allocation strategy of the cell, that is, the interplay between cellular functions such as growth, nutrient uptake, respiration, and photosynthesis (Toseland *et al*., 2013; Zimmerman *et al*., 2014; Hui *et al*., 2015; Inomura *et al*., 2020). In turn, the changes in resource allocation can affect the elemental and macromolecular composition of the cell.

All living organisms are composed of a number of chemical elements, including macroelements, (i.e., C, N and P) and microelements, i.e., trace metals such as iron (Basu *et al*., 2015). These elements are biochemically combined as part of the macromolecular building blocks that comprise most of the cell biomass - carbohydrates, proteins, nucleic acids, lipids and a vast array of various metabolites. Of these pools, proteins are typically the largest in terms of dry weight, followed by carbohydrates and nucleic acids or phospholipids (Table S1). Most of the C in the cells is found in carbohydrates, while N is mostly found as part of proteins (Table S2, (Moreno and Martiny, 2018)). It has been suggested that cellular P is associated mainly with RNA molecules, and to a lesser extent DNA, phospholipids and “P-esters” (e.g. phosphorylated proteins and metabolites) (Sterner and Elser, 2002; Rees and Raven, 2021). However, there are other pools of P not associated with these specific macromolecules, for example polyphosphate stores, which are often associated with ‘luxury uptake’ in response to nutrient pulses under P-limited conditions (Kulaev et al., 2005). Phosphonates, organophosphorus metabolites, are another potential storage of P that may be used for secondary metabolism (Acker *et al*., 2022). Thus, in order to understand the elemental composition of marine organic matter, there is a need to study the environmental conditions affecting the intracellular resource allocation of macromolecules.

Temperature is one of the environmental conditions that strongly affects the growth rate, physiology, and macromolecular composition of bacterioplankton. Two different hypotheses have been raised as to how temperature affects resource allocation in microorganisms. According to the translation-compensation hypothesis (TCH, Fig 1A), the main factor affecting the cell requirement for ribosomes (and thus the amount of cellular RNA) is the biophysical effect of temperature on enzymatic activity rates (Fraser *et al*., 2002; Toseland *et al*., 2013). Where, at low temperatures cells require more ribosomes to compensate for the lower rate of translation, while an increase in temperature increases translation efficiency and reduces the number of required ribosomes. Evidence supporting the TCH comes from multiple bacteria (e.g. *E. coli, Pseudomonas fluorescens* and others) grown under nutrient-replete conditions, where cellular P and/or RNA content increase with decreasing temperatures (Woods *et al*., 2003; Cotner *et al*., 2006; Chrzanowski and Grover, 2008). It has also been suggested that the decrease in N:P ratio observed in environmental samples with decreasing temperature at higher latitudes which results in non-Redfield ratio (Redfield, 1934, 1958), is driven by the increased need for ribosomes, also supporting the TCH (Daines et al., 2014; Toseland et al., 2013; Yvon-Durocher et al., 2015).

**Fig. 1.**
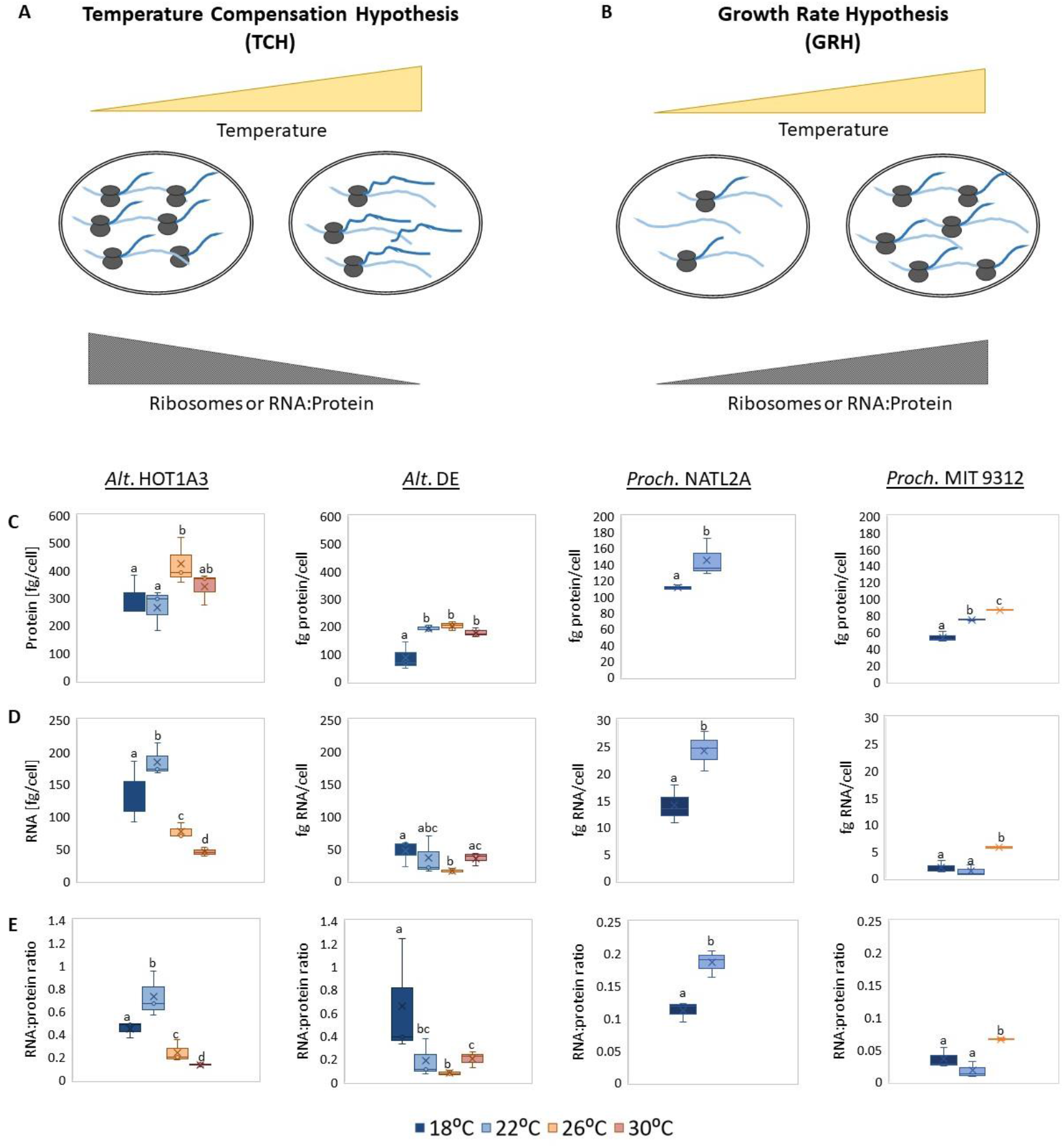
Testing the TCH and GRH in laboratory cultures of *Alteromonas* and *Prochlorococcus*. (A, B) Schematic illustrations of the TCH (A) and GRH (B). The blue lines emerging from the ribosome-RNA complexes are the nascent proteins, which according to the TCH are longer at higher temperatures due to the higher processivity of the ribosomes. C, D) Temperature effect on *Alteromonas* and *Prochlorococcus* protein (C) and RNA (D) per-cell quotas. Concentrations were normalized to cell counts obtained by flow cytometry. E) RNA:protein ratio. Note the different Y-axis between *Alteromonas* and *Prochlorococcus*. Box plots show the average and standard deviation of 3 biological replicates. The different letters above the box plots in the graphs indicate statistically significant differences among the test temperatures (one-way ANOVA, *P*<0.05). *Prochlorococcus* MIT9312 and NATL2A did not grow at 30°C and 26-30°C respectively and therefore no information is available.

An alternative hypothesis, the growth rate hypothesis (GRH), states that growth rate depends primarily on the availability of P-rich ribosomes for protein production (Fig 1B, Sterner & Elser, 2002). Increased RNA content with growth rate was observed for example in *E. coli* (Hanegraaf and Muller, 2001; Makino *et al*., 2003) and in different isolates of marine bacteria (Kemp *et al*., 1993). Both P and RNA content also increased with temperature in freshwater bacteria, particularly under P-limitation (Phillips *et al*., 2017). Thus, while temperature can strongly affect the resource allocation strategy of the cells (the interplay between protein and ribosome synthesis), the exact effect of temperature on resource allocation and thus elemental ratios remains unclear, as can be seen by the opposite predictions of the TCH and the GRH.

Here, we examined which of the two hypotheses, the TCH or the GRH, best explains the macromolecular composition of the ecologically-important ubiquitous marine cyanobacterium *Prochlorococcus* and the heterotrophic marine bacterium *Alteromonas. Prochlorococcus* is the smallest and most abundant cyanobacteria in the ocean (reviewed by (Partensky and Garczarek, 2010; Biller *et al*., 2015)). *Alteromonas* is a clade of heterotrophic marine bacteria that tend to dominate 16S libraries at certain locations and times (Acinas et al., 1999; Garcia-Martinez et al., 2002; Roth Rosenberg et al., 2021). Additionally, we characterized how the protein and RNA quotas of natural bacterioplankton communities changed over time in the nutrient-poor Eastern Mediterranean Sea, aiming to identify which environmental factors were correlated with the RNA:protein ratio.

### Experimental Procedures

#### Culturing Alteromonas and Prochlorococcus

Cultures of *Alteromonas* and *Prochlorococcus* were grown under different temperatures to assess if and how temperature affects their macromolecular composition. Triplicate *Alteromonas* cultures strains were grown in ProMM (Morris *et al*., 2008) for 11h after 48h of acclimation to the growth temperature. Cultures of *Prochlorococcus* were grown under continuous light (∼20µMole Quanta/m^2^×sec) and maintained in Pro99 (Moore et al., 2007). Each strain was acclimated to the experimental temperature for at least five transfers before the experiment was started by inoculating an initial density of ∼2 ×10^−6^ cells/ml.

#### Sampling procedure

For *Alteromonas*, culture optical density was measured every 1h, and for *Prochlorococcus*, fluorescence was measured every 1-3 days. To isolate the effect of temperature from both growth and nutrient starvation, cultures were sampled at mid-exponential growth, which was identified in several preliminary experiments. Samples (10-20mL) were collected for measurements of particulate organic P (POP), protein, RNA, polyphosphates, and cells count. Protein, RNA and polyphosphates samples were preserved in lysis buffer (40mM EDTA, 50mM Tris pH=8.3, 0.75M sucrose), frozen and stored at −80°C. Particulate P samples were frozen and stored at −20°C. Samples for flow cytometry were fixed with 25% glutaraldehyde for ten minutes, frozen and stored at −80°C.

#### Monthly field sampling in the Eastern Mediterranean Sea

Seawater samples were collected onboard the R/V MedEx or R/V Bat-Galim from the surface (2.5 m) and the Deep Chlorophyll Maximum (DCM) (∼50-135 m) at the THEMO2 offshore station, located ∼30 miles west of Haifa (∼1450 m bottom depth, Lat. 32° 47.49, Lon. 34° 22.50). Samples were also collected from 500 m depth, but the concentrations of RNA and protein were below the limit of detection and are therefore not discussed further. Eight liters of seawater were filtered for the following measurements: particulate P (duplicates), protein and RNA concentration (triplicates or quadruplicates). Samples were collected and preserved as described above (protein and RNA samples were first frozen in liquid nitrogen and then stored at −80°C). In addition, complementary measurements were taken during the cruises. This includes a full Conductivity Temperature and Depth (CTD) profile, algal fluorescence, oxygen levels, picophytoplankton and heterotrophic cells count, bacterial production and primary production (Reich et al., 2021) and dissolved nutrients (NOx, PO4, (Ben Ezra *et al*., 2021)).

#### Sampling in the South China Sea

Seawater samples were collected on cruise HKB-SCS-2021 from the surface (2-5 m) and DCM layer (5-70 m) at 3 open water stations and 5 stations near the Estuary (Fig S7). Between 1-6 liters of seawater were filtered for protein (1 replicate) and RNA (duplicates). Samples were collected and preserved as described above and shipped to Israel on dry ice. POP was measured on particles on 0.2 *µ*m polycarbonate membrane filters using the persulfate oxidation method (Menzel and Corwin, 1965).

#### Measurements of macromolecules and particulate P

Proteins were extracted by incubating with lysis buffer for 30 min at 37°C followed by 10 min sonication in a water bath (Ultrasonic cleaner AC-120H) and measured using the Bicinchoninic Acid (BCA) method (Smith *et al*., 1985). RNA isolation from the filter was based on Trizol extraction (Chomczynski and Sacchi, 1987). RNA concentration was measured using Qubit® RNA HS Assay Kit. POP samples were filtered on pre-combusted (550°C, 2h) 47mm (8L, field samples) or 25mm (10mL, laboratory cultures) glass fiber filters (GF/F, Whatman) and stored at −20°C until further analysis. Samples were dried in an oven (60°C overnight) prior to analysis using the persulfate oxidation method (Menzel and Corwin, 1965). P quotas of polyphosphates and POP were calculated by dividing the measured PO_4_ concentrations by 0.3261 (based on PO_4_ molecular weight), and P-RNA was calculated as 9.1% of the RNA mass (Geider and La Roche, 2002).

#### Measurements of polyphosphates

Polyphosphate extraction was done using an adjusted phenol-chloroform extraction (Bru *et al*., 2017). Concentrations were multiple by 1.4 to account for ∼40% loss of polyphosphate during the extraction (Christ and Blank, 2018). Concentrations of polyphosphate were measured using the Phosfinity Total Polyphosphate Quantification Kit.

#### Estimation of P in DNA and lipids

P-DNA was calculated from the estimated total P atoms in the genome of each bacterium (one P atom per nucleotide x total nucleotides), assuming 1.5 chromosomal copies to account for DNA replication during exponential growth. P-phospholipids was estimated based on the surface area of the cells, taking into account (for *Prochlorococcus*) also the thylakoids, according to Waldbauer *et al*. (2012). We assumed a spherical cell 0.6µM in diameter for *Prochlorococcus* and a rod-shaped cell 0.6×1.8µM for *Alteromonas* (based on preliminary SEM images taken during the exponential growth). P-phospholipids (1 fg/cell for *Alteromonas* and 0.1 and 0.065 for NATL2A and MIT9312) was estimated according to the percent of phospholipids out of the total lipids: 85% for the heterotrophic *Alteromonas*, 14% for NATL2A and 9% for MIT9312 (Van Mooy *et al*., 2006). Estimations for *A. macleodii* HOT1A3 were multiplied by 3 to account for the overestimation of cellular quotas (P, RNA and polyphosphates) following the difficulties in cell counting (see SI and Fig S4).

## Results and Discussion

### Differences in macromolecules quotas between Alteromonas and Prochlorococcus strains

Before assessing the effect of temperature on macromolecular composition in *Alteromonas*, preliminary experiments were performed in order to determine how growth rates depend on temperature in diverse isolates (Fig S1). Growth rates increased monotonously from 18°C to 30°C in all isolates, decreasing at 32°C in *A. macleodii* HOT1A3 but not in other strains (*A. macleodii* ATCC and BS11 and *A. mediterranea* DE and DE1). There were notable differences in growth rate and in the shape of the growth curve observed between *A. macleodii* and *A. mediterranea* strains, which are discussed in the Supporting information and Fig S1. Based on these results, we selected two strains of *Alteromonas, A. macleodii* HOT1A3 (a ‘surface ecotype’) and *A. mediterranea* DE (a ‘deep ecotype’), for further analysis at larger culture volumes between 18°C and 30°C (Fig S2). Moreover, two strains of *Prochlorococcus* were selected, the high-light (HL) adapted MIT9312 (a ‘surface ecotype’) and the low-light (LL) adapted NATL2A (a ‘deep ecotype’), as they have different temperature optima (Zinser *et al*., 2007) that undergo a successive pattern over time (Malmstrom *et al*., 2010). Experiments with *Prochlorococcus* were performed at 18°C-26°C for MIT9312 (whose permissive range is up to ∼28°C, (Ma *et al*., 2018)) and 18°C-22°C for NATL2A, which failed to grow at higher temperatures (Fig S3).

Overall, per-cell quotas of protein and RNA were all highest for *A. macleodii* HOT1A3, followed by *A. mediterranea* DE, *Prochlorococcus* NATL2A, and finally *Prochlorococcus* MIT9312 (Fig 1C and 1D). The larger, copiotrophic *Alteromonas* is expected to have higher cell quotas of macromolecules compared to *Prochlorococcus* which is often associated with oligotrophic environments, due to their size and a larger number of ribosomal RNA operons (5 in *Alteromonas* compared to 1-2 in *Prochlorococcus*) (Rocap *et al*., 2002; Schirrmeister *et al*., 2012; López-Pérez *et al*., 2013; Rees and Raven, 2021). Yet, the per-cell protein quotas of *A. macleodii* HOT1A3 (∼200-500fg/cell) were considerably higher than those of *A. mediterranea* DE (50-200fg/cell). The high values are within the range of other marine copiotrophic bacteria such as *Vibrio splendidus* assuming proteins are 50% of the dry cell mass (Cermak *et al*., 2017), whereas the low ranges are similar to a different *Alteromonas* strain, Alt1C (70 fg/cell protein and ∼14 fg/cell RNA, (Zimmerman *et al*., 2014)). The reasons for the differences in quota between the *Alteromonas* strains are currently unclear but may be related to their different tendencies to form floating biofilms, which can affect the estimation of cell number by flow cytometry (see SI and Fig S4 for a comparison of biofilm formation between *Alteromonas* strains). The per-cell protein quotas for *Prochlorococcus* are consistent with or slightly higher than previous studies: ∼60-140 fg/cell, compared with ∼30-130 fg (Cermak *et al*., 2017; Roth-rosenberg *et al*., 2020; Casey *et al*., 2022). The higher values for *Prochlorococcus* NATL2A compared to *Prochlorococcus* MIT9312 are in agreement with the difference in size between the two strains (Cermak *et al*., 2017). The RNA quotas for MIT9312 are somewhat lower than previously reported (∼1-6 fg/cell compared with ∼9fg/cell for MIT9312 (Roth-rosenberg *et al*., 2020) and ∼3-4fg/cell for MED4 (Casey *et al*., 2022)).

### Temperature affects resource allocation differently in Alteromonas and Prochlorococcus

The changes in temperature affected differently the RNA:protein ratio in *Alteromonas* and *Prochlorococcus* (Fig 1E). Broadly speaking, a tendency was observed for the RNA:protein ratio of *Alteromonas* to decrease with temperature, in accordance with the TCH. In contrast, the RNA:protein ratio increased with temperature in *Prochlorococcus*, as predicted by the GRH. In strain NATL2A, the RNA:protein ratio also increased with culture growth rate, further supporting the GRH in this strain, whereas it decreased in *Alteromonas* HOT1A3 (Fig S5). These changes were mostly determined by the changes in RNA quotas rather than the protein (Fig 1D). The decrease in RNA:protein with temperature, as observed in *Alteromonas* and predicted by the TCH, is in accordance with observations in other bacteria (Woods *et al*., 2003; Chrzanowski and Grover, 2008) and diatoms (Toseland *et al*., 2013). Similarly, the increase in RNA:protein in *Prochlorococcus* is in accordance with a previous study that observed an increased P:N ratio with temperature in *Prochlorococcus* strain MIT9312 (Martiny et al., 2016) as well as in freshwater bacteria (Phillips *et al*., 2017).

On average, the RNA:protein ratios of *Alteromonas* were ∼2-10 fold higher compared to *Prochlorococcus* (Fig 1E), only partly consistent with the ∼20-fold higher growth rates (Fig S2 and S3). One possible explanation for this is that ribosomes in *Prochlorococcus* may work closer to their maximal capacity, as they need to support also the needs of the photosynthetic apparatus. Nonetheless, the RNA:protein ratio of NATL2A was more similar to *Alteromonas*, while that of MIT9312 was considerably lower. The RNA:protein ratios here are broadly consistent with other studies, where it was ∼0.03-0.08 for MIT9312, ∼0.08 for MED4 and ∼0.2-0.6 for heterotrophs (Roth-rosenberg *et al*., 2020; Casey *et al*., 2022).

Notably, in none of the cases we observed a monotonic increase or a decrease in the RNA:protein ratios, which would have been in strict accordance with the GRH or the TCH. For example, for *A. macleodii* HOT1A3 the highest RNA:protein ratio was observed at the intermediate temperature of 22°C, and a slight but significant increase was observed at the highest temperature in *A. mediterranea* DE (Fig 1E). This can be due to a shift in growth and resource allocation strategy, where the GRH and the TCH each apply at different temperature ranges. For example, an increase in RNA quotas at the highest temperature in *E. coli* (as in *A. mediterranea* DE) was previously explained by increase in the ribosomes catalytic capacity at temperatures above those optimal for growth (e.g. due to increased protein turnover as a result of misfolding (Farewell and Neidhardt, 1998; Cotner *et al*., 2006)).

### Ribosomes are not always the main P reservoir

In order to link the cell’s macromolecular composition (protein and RNA) and elemental (N:P) ratio, it is commonly assumed that most of the P in the cell is invested in RNA (Sterner and Elser, 2002). To test this, we measured total cellular P quota (particulate organic P divided by cell number, Fig 2A) and calculated the fraction of the P that is allocated to RNA (P-RNA:P) (Fig S6A). Overall, the measured P quotas for *Prochlorococcus* (2-2.5fg/cell) were 2-5 fold higher than previously reported (∼0.5-1fg/cell (Bertilsson et al., 2003; Casey et al., 2022; Martiny et al., 2016). In contrast, the cellular P quotas of *A. mediterranea* DE are similar to those reported for Alt1C (5.8 fg P/cell, (Zimmerman *et al*., 2014)). P-RNA:P and RNA:protein were correlated in all tested organisms (Fig S5C). However, despite this correlation, there was a very wide range of P-RNA:P values, and in many cases only a relatively small fraction of the cellular P (as small as 8%) was associated with RNA (Fig 2B, Fig S6A). This wide range is observed also in a literature compilation across bacteria (Table S3).

**Fig. 2.**
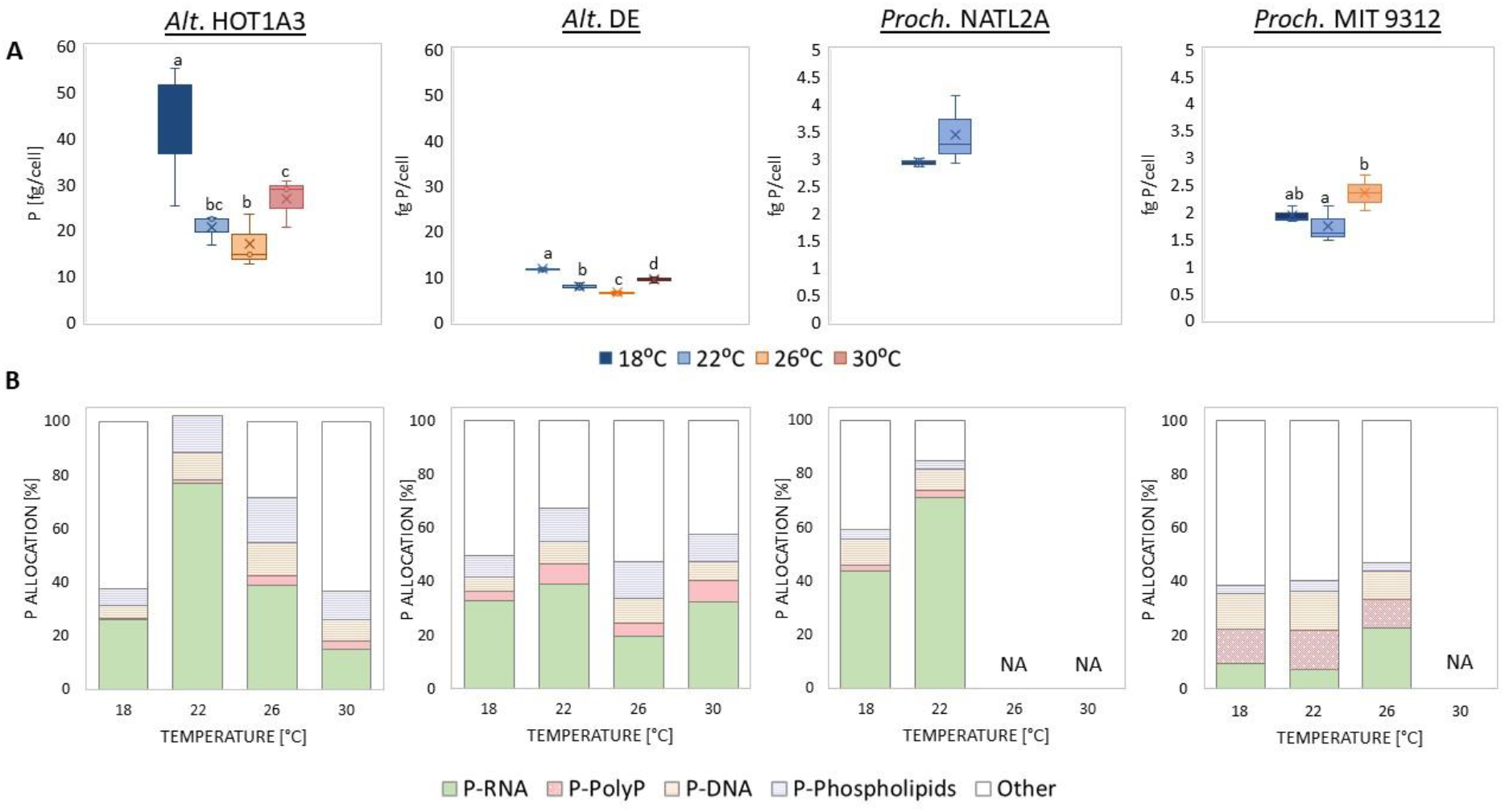
Temperature effect on *Alteromonas* and *Prochlorococcus* phosphorous allocation. (A) Total cellular phosphorus quotas. Boxplots show average and 75 percentile from three replicate cultures. The different letters above the box plots in the graphs indicate statistically significant differences among the test temperatures (one-way ANOVA, *P*<0.05). (B) Allocation of P to RNA, polyphosphates (PolyP), DNA, and phospholipids. Concentrations of P, RNA and polyphosphates were measured, while those of DNA and phospholipids were estimated as described in the materials and methods. Measurements of polyphosphates for MIT9312 were taken from a different experiment at 26°C since the concentrations of the original samples were below the limit of detection (<5µM). NA= not available, since the cells could not grow in these temperatures.

P-RNA:P varied between the bacteria and, in some cases, also between temperatures. In *A. macleodii* HOT1A3, P-RNA:P was higher in the middle of the temperature range compared to the edges (∼40-80% compared to 15-30%, Fig S6A), while in *A. mediterranea* DE P-RNA:P was ∼20-40% across the entire temperature range. These values are somewhat higher than those measured in Alt1C, (∼20%, (Zimmerman *et al*., 2014)) yet within the range of previous studies in *E. coli* (48-62%, (Cotner *et al*., 2006)). Both *Prochlorococcus* strains exhibited a significant increase in P-RNA:P with temperature (Fig S6A). However, while P-RNA:P in NATL2A was similar to *Alteromonas*, that of MIT9312 was lower, more in line with previous observations in *Prochlorococcus* MED4 (20-30%, (Casey *et al*., 2022)) and *Synechococcus sp*. WH8102 (5-15% (Mouginot *et al*., 2015; Garcia *et al*., 2016)).

The lower P-RNA:P observed in MIT9312 and in some cases in *Alteromonas* prompted us to measure or calculate other potential P stores. Therefore, we measured polyphosphates, while P-DNA was estimated based on the genome size of each bacterium and P-phospholipids was estimated from the total lipids in the cell membrane (Waldbauer *et al*., 2012) (Fig 2B). The calculated fractions of P in DNA (8-13%) and phospholipids (3-12%) were in the same range as reported for heterotrophic bacteria and cyanobacteria (see Table S2 and (Rhee, 1973; Cotner *et al*., 2006; Rees and Raven, 2021). In contrast, the fraction of P in polyphosphates was only 1-15% in both *Alteromonas* and *Prochlorococcus* (Fig 2B, Fig S6B), lower than previously suggested in cyanobacteria (29±9%, (Rees and Raven, 2021) and Table S2). However, the polyphosphates quotas measured here are in accordance with a previous study in MED4 (0.25-0.3 fg/cell), although in that study the total cellular P quota was lower, resulting in a higher relative contribution of polyphosphates (25-30% of cellular P (Casey *et al*., 2022)). It is to be noted, though, that polyphosphates estimates in prokaryotes are scarce, highlighting the need for more measurements of this P storage pool.

The cellular P budget detailed above suggests that a considerable fraction of cellular P in all tested strains, and under most growth conditions, is not found in one of the measured or calculated macromolecular pools. Several hypotheses may account for this “missing P”, which can reach as much as 60% of the total cell quota. Firstly, we may have underestimated the amount of polyphosphates or RNA due to loss of material during the extraction process, or overestimated the cellular P quotas, although the latter option is less likely as P estimates from field samples were similar to previous studies (see below). Secondly, part of the unaccounted-for P may be associated with phosphate-esters and anhydrides (e.g. phosphorylated proteins and metabolites such as ATP (Rees and Raven, 2021)). Thirdly, a recent study suggested that phosphonates may also be an important reservoir of P in *Prochlorococcu*s, and may account for up to 40% of the cellular P in some strains (Acker *et al*., 2022). However, while the two *Prochlorococcus* strains tested here contain the phosphonate transporter gene *phnD* (Ilikchyan *et al*., 2009), phosphonatase or C-P lyase that are needed for phosphonate utilization were not identified in their genome (Martiny *et al*., 2006; Acker *et al*., 2022).

### The macromolecular quotas between axenic lab cultures and mixed field bacterial communities considerably differ

To what extent are the macromolecular quotas from axenic laboratory cultures representative of natural microbial communities? To test this, we collected samples in a monthly time-series in the Eastern Mediterranean (surface and Deep Chlorophyll Maximum, DCM), as part of the SoMMoS project (Table 1, Fig S7, (Reich et al., 2021)). Overall, protein quotas per cell were somewhat lower in the field compared to cultures of *Alteromonas* and *Prochlorococcus*, being closest to those of *Prochlorococcus* MIT9312 (Table 1). These values are within the range observed in natural communities, e.g. from the Scripps Pier (∼20-60 fg/cell, (Simon and Azam, 1989)) and from the estuarine and coastal waters in the Northern Gulf of Mexico (∼29±12 fg/cell, (Jeffrey *et al*., 1996)). Cellular P quotas were typically ∼2-fold lower than *Prochlorococcus* MIT9312 and ∼10-fold lower than *A. mediterranea* DE. RNA quotas, however, were significantly lower in the field compared with laboratory cultures: ∼3-100 fold lower than *Prochlorococcus* MIT9312 and up to 7000-fold lower than *A. mediterranea* DE (Table 1). This results also in very low RNA:protein and P-RNA:P ratios, up to ∼1,000-fold lower than in lab cultures. Such very low RNA:protein ratios (and P-RNA:P) were also observed in samples from a transect from the Pearl River Estuary to the South China Sea (Fig S8E, F), with higher ratios in the Pearl River Estuary, similar to MIT9312 and close to the lower range of NATL2A and *A. mediterranea* DE (Table 1).

**Table 1.** Results of the physiological mmeasurements of laboratory cultures of *Alteromonas* and *Prochlorococcus* compared to the field (*in-situ*). Values shown are ranges (minimum and maximum).

|  | *Alt* HOT1A3 | *Alt* DE | NATL2A | MIT9312 | E. Med Surface | E. Med DCM | South China Sea | Pearl River Estuary |
| --- | --- | --- | --- | --- | --- | --- | --- | --- |
| Protein [fg/cell] | 181-520 | 47-218 | 109-172 | 50-87 | 6-85 | 13.5-36 | NA | NA |
| P [fg/cell] | 14-62 | 7-13 | 2.8-4.2 | 1.5-2.7 | 0.03-3.6 | 0.26-2.2 | NA | NA |
| RNA [fg/cell] | 40-214 | 14-70 | 11-28 | 1-6 | 0.02-0.35 | 0.01-0.25 | NA | NA |
| RNA:protein ratio | 0.12-0.95 | 0.06-1.25 | 0.094-0.21 | 0.008-0.07 | 0.0008-0.013 | 0.0006-0.01 | 0.0001-0.0027 | 0.002-0.057 |
| P-RNA:P [%] | 13-87 | 16-71 | 34-78 | 3-27 | 0.2-4.1 | 0.15-1.83 | 0.3-11 | 26-32 |

*Alteromonas* HOT1A3 and DE data include all measurements from 18°C, 22°C, 26°C and 30°C (N=12). *Prochlorococcus* MIT9312 includes all measurements from 18°C, 22°C and 26°C (N=9), and NATL2A all measurements from 18°C and 22°C (N=6). Data from the Eastern Mediterranean include all measurements for POP (N=11), months Jun-Jan for protein and RNA (N=7) and months May-Sep for PON (N=5). Data from the South China Sea include 3 stations (including a station that represents the transitional zone), and that of Pearl River Estuary 5 stations. NT= not tested. NA=not available.

The lower P quotas observed in natural populations compared to laboratory cultures are unlikely to be due to analytical biases in field samples, as the measurements of particulate organic phosphorus and of cell counts are in the same range as previous studies from the Levantine Basin (Tanaka *et al*., 2011). Measurements of RNA/cell in natural communities are scarce, making it more difficult to evaluate the measurements from the Eastern Mediterranean. Simon and Azam failed to measure RNA in pelagic bacteria, which they explained by either very low RNA content (<1 fg/cell) or RNA degradation during sample preparation (Simon and Azam, 1989). Similarly, a study of natural bacterioplankton samples from a coastal region of Long Island suggested that up to ∼78% of the cells during summer had less than 0.3 fg RNA/cell (Lee & Kemp, 1994). Notably, in cells where RNA could be measured by Lee and Kemp (1994), the quota was 1.6-5.4 fg/cell, within the range of *Prochlorococcus* MIT9312 cultures (Table 1). Therefore, while additional field studies are needed to verify our results, it seems that the average RNA/cell is very low in natural communities.

We propose four, non-exclusive, hypotheses, to explain the much lower P and RNA quotas observed in natural, oligotrophic communities. Firstly, the natural microbial community in oligotrophic oceans (including the Eastern Mediterranean) is numerically dominated by small, oligotrophic bacteria such as SAR11 (Giovannoni, 2017; Roth-Rosenberg et al., 2021). The dry mass of laboratory-cultured SAR11 strains is 5-10 fold lower than *Prochlorococcus* (Cermak *et al*., 2017), consistent with the ∼10-fold lower P quotas in natural SAR11-dominated communities. Secondly, previous studies have shown that cells from laboratory cultures of *Prochlorococcus* are smaller than those of field populations, and have fewer ribosomes per cell (e.g. (Lin *et al*., 2013)). Similarly, a proxy for cell size using flow cytometry (based on forward scatter) showed that *Prochlorococcus* in the lab are ∼3 fold larger than in the field (Fig S9). Thirdly, it is possible that a significant fraction of the cells in natural communities are non-active or non-living (e.g. (Lasternas *et al*., 2010)). Finally, it is possible that other macromolecular pools, such as polyphosphates or phospholipids, constitute most of the P quotas in cells exposed to natural, nutrient limited, conditions (Daines *et al*., 2014).

### Testing the TCH and the GRH in the Eastern Mediterranean Sea

We next asked whether the TCH and/or the GRH are able to explain the variation in the macromolecular composition of the natural community in the Eastern Mediterranean over time. Sea surface temperature changed from ∼18°C during winter to ∼29°C during summer (Reich et al., 2021), and there was a negative correlation between the surface temperature and both surface RNA:protein and P-RNA:P (Fig 3A). This suggests that as temperature increased the allocation of resources to ribosomes by surface microbial communities also increased, in contrast with the prediction of the TCH (Toseland et al., 2013). In contrast to the surface water, the temperature in the DCM remained constant throughout the year (17.7±0.39°C, (Reich et al., 2021)). Therefore, while changes were observed in the RNA:protein and P-RNA:P ratios in the DCM (Fig S6E and S6F), these too cannot be explained by the TCH.

**Fig. 3.**
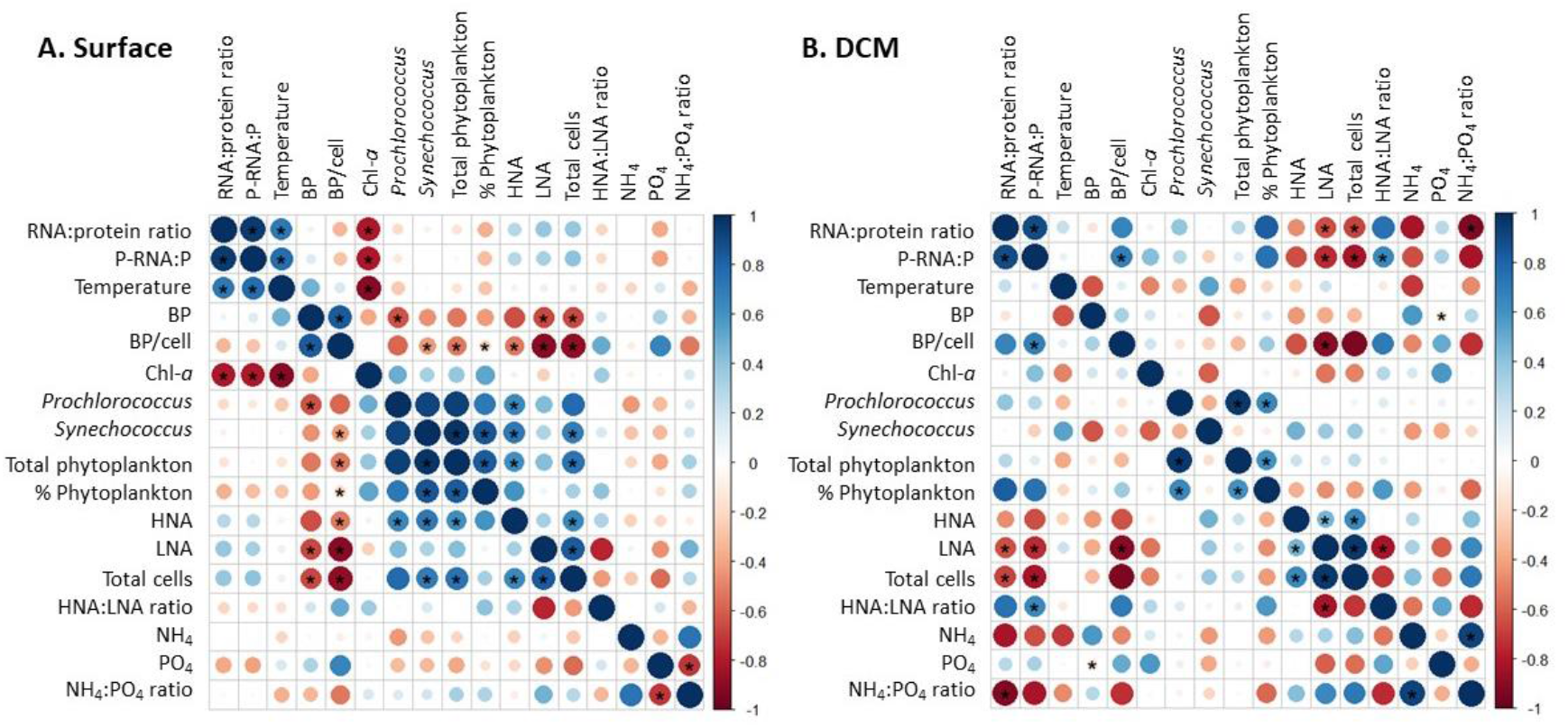
Matrix of Spearman’s correlation coefficients between all environmental variables. (A) Surface water. (B) Deep Chlorophyll maximum (DCM) layer. List of variables by order: RNA:protein ratio; P-RNA:P [%]; Temperature [°C]; Bacterial production [BP] [ng C L^-3^ h^-1^]; BP/cell [fg C h^-1^]; Chlorophyll-*a* [µg/L]; *Prochlorococcus* [cells/mL]; *Synechococcus* [cells/mL]; Total phytoplankton [cells/mL]; Total phytoplankton [%]; High nucleic acids [HNA] [cells/mL]; Low nucleic acids [LNA] [cells/mL]; Total cells [cells/mL]; HNA:LNA ratio; NH_4_ [µM]; PO_4_ [µM]; NH_4_:PO_4_ ratio. Points are coloured and sized according to the value of the Spearman’s correlation coefficient (blue dots represent positive correlation and red dotes represent negative). Asterisks indicating the significance at *p* < 0.05 level. Plot created using the corrplot package in R (Wei et al., 2017).

Unlike the TCH, which can be tested by correlating macromolecular composition with the easily measurable temperature, testing the GRH in the field requires an estimate of community growth rate. We chose to use per-cell bacterial production (BP), which is widely employed to assess growth rates of bacteria in the oceans (Kirchman et al., 1985), despite some limitation such as the assumption of constant cell size (e.g. (Kirchman, 2016)). Assuming an average cellular C quota of 20 fg C/cell (Lee & Fuhrman, 1987), the average growth rate (based on the BP presented by Reich et al., 2021) was 0.35±0.26 d^-1^ (range 0.1-1 d^-1^). This is within the range of published growth rates for *Prochlorococcus* measured in the lab and *in-situ* using multiple approaches (0.3-1d^-1^, e.g. (Vaulot *et al*., 1995; Hunter-cevera *et al*., 2014; Zubkov, 2014; Ribalet *et al*., 2015)) and somewhat higher than lab cultures of SAR11 (0.216 d^-1^ (Zimmerman *et al*., 2014)). It is also similar to estimates from other oceanic regions (Kirchman, 2016). Per-cell BP was positively and significantly correlated with P-RNA:P in the DCM but not at the surface (r=0.6258, *P*-value=0.007, Fig 3). Per-cell BP was also positively correlated with RNA:protein, although this correlation was not statistically significant (r=0.5128, *P*-value=0.076). Thus, there is limited support for the GRH in the DCM but not at the surface.

In addition to temperature and growth rate, other factors may affect the community RNA:protein and P-RNA:P. At the surface, RNA:protein and P-RNA:P were negatively correlated with chlorophyll-*a* (Fig 3A), whereas at the DCM they were negatively correlated with the total number of cells as well as the number of low nucleic acid (LNA) cells, and P-RNA:P was also positively correlated with BP/cell and HNA:LNA ratio (Fig 3B). This suggests a general trend for lower RNA:protein under oligotrophic conditions when phytoplankton biomass is low and there are many small or less active cells. Finally, RNA:protein was also negatively correlated with the N:P ratio of inorganic nutrients. The Eastern Mediterranean is often considered to be P and/or N and P co-limited (Zohary *et al*., 2005; Tanaka *et al*., 2011) and these results suggest that the relative availability of N and P might determine cellular resource allocation.

### Conserved variation in resource allocation between the axenic lab cultures and mixed field bacterial communities

As described above, there are clear differences between the lab cultures and the *in-situ* microbial populations of the Eastern Mediterranean Sea, both in terms of RNA quotas and P allocation, and in the applicability of the TCH or GRH. Nevertheless, there is a consistent relationship across laboratory and field samples in the ratio between RNA and P quotas (Fig 4A) and in the ratio between RNA:protein and P-RNA:P (Fig 4B), which both follow power laws. Cells with higher P quotas also have higher amounts of RNA, and likely more ribosomes (Fig 4A). The correlation, though, is weaker when only field samples are considered. It suggests that about ∼40% of the variability in P quotas can be explained by the changes in RNA, while the remaining ∼60% is due to variable cellular quotas of other P-containing macromolecules. The correlation between the RNA:Protein and P-RNA:P (Fig 4B) suggests that, even though RNA may be a relatively minor reservoir of cellular P, much of the variability in P allocation between different cellular pools is nevertheless related to the cellular need to maintain sufficient ribosomes for protein production and turnover. Here, there is more overlap between the field and lab samples, especially *Prochlorococcus* (Fig 4B, blue trendline). Overall, these correlations suggest that the regulation of resource allocation and life strategy of the oligotrophic and copiotrophic bacteria in the lab is not considerably different from that of natural communities of bacteria in the sea.

**Fig. 4.**
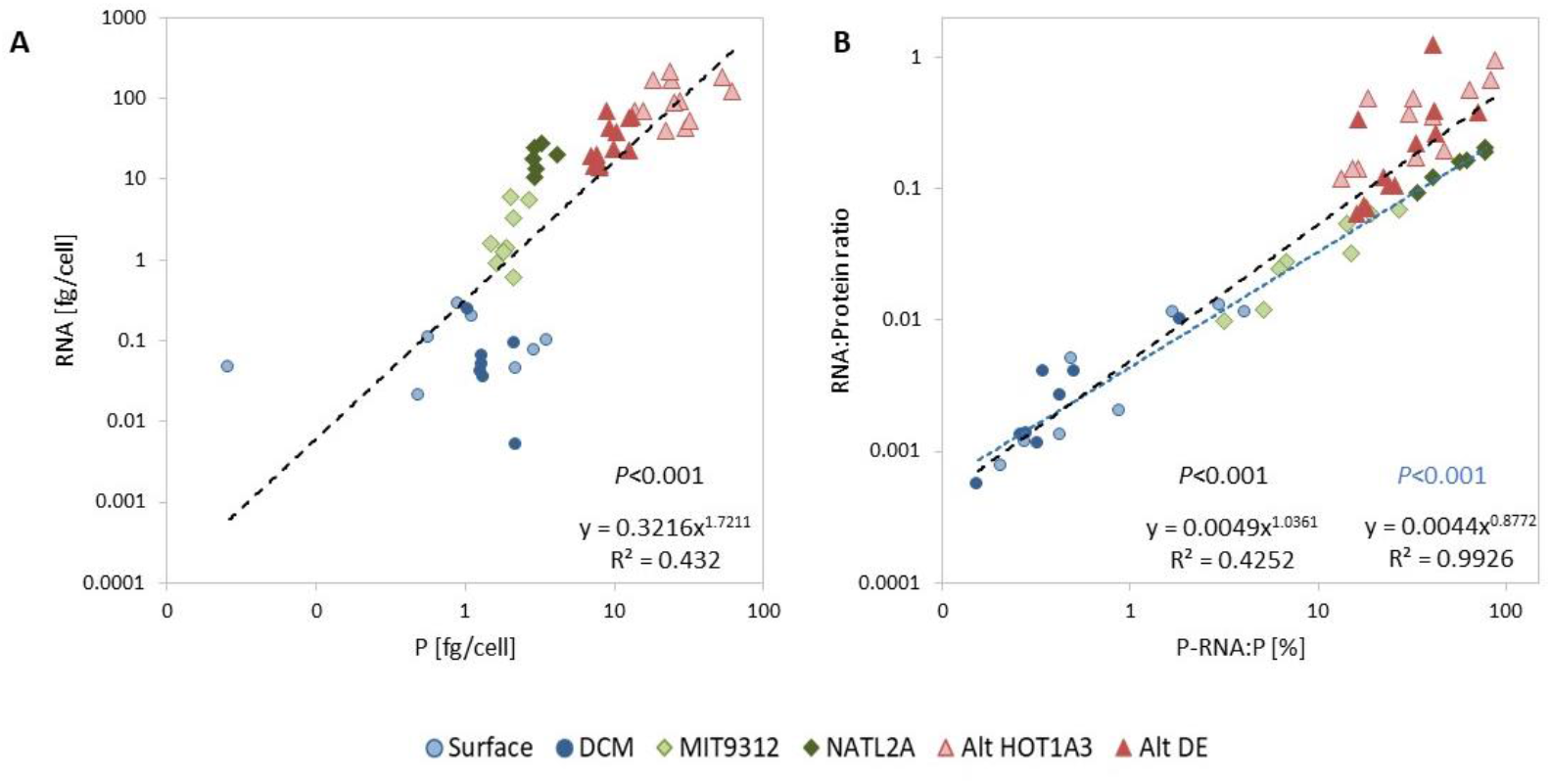
Correlation between cellular quotas of RNA and P (A) and between the allocation of P to RNA and the RNA:protein ratio (B). The black trend lines show power law correlations, the blue line in panel B shows the same correlation for only field and *Prochlorococcus* samples (without *Alteromonas*). Note that logarithmic axes. *Alteromonas* HOT1A3 and DE data include all measurements from 18°C, 22°C, 26°C and 30°C (N=12). *Prochlorococcus* MIT9312 includes all measurements from 18°C, 22°C and 26°C (N=9), and NATL2A all measurements from 18°C and 22°C (N=6). Eastern Mediterranean data includes all measurements for POP (N=11) and measurements from June 2018 to January 2019 for protein and RNA (Fig S6, N=7).

### Summary and future prospects

In this study, we aimed to test the TCH and GRH *in-vitro* with two very different clades of bacteria (an oligotrophic primary producer and a copiotrophic heterotroph), as well as *in-situ*. Support for the TCH was found only in the copiotrophic *Alteromonas*, while the result of the oligotrophic *Prochlorococcus*, and to a lesser degree natural samples from the DCM, provide some support for the GRH. However, the cellular response to changes in temperature was not a simple, monotonic change in RNA:protein ratios. Rather, more complex patterns were observed, suggesting that aspects of both hypotheses are acting together, for example, non-linear increases in both ribosome efficiency and ribosome requirements with temperature. Further studies, potentially using chemostats to better control growth rate and detailed analyses of the rates of protein production and degradation, are needed in order to dissect the effect of these contrasting mechanisms. It is also possible that the GRH is best formulated in terms of maximal rather than actual (instantaneous) growth rate, and that molecular-genomic aspects such as the number of rRNA operons are also important (Rees and Raven, 2021).

A considerable amount of P was not allocated to RNA, especially in MIT9312 and in the field samples. Previous studies have observed strong latitudinal and seasonal patterns in N:P ratios, and in some cases this has been explained through the TCH and GRH (Martiny *et al*., 2013; Toseland *et al*., 2013; Yvon-Durocher *et al*., 2015, 2017). The lack of support for either of these hypotheses in the Eastern Mediterranean, and the limited coupling between P and RNA quotas, urge caution in interpreting the variability in elemental ratios in light of these two hypotheses. These results also call for a more detailed analysis of the various P pools in marine microbes, as these are likely to play a large role in oceanic particulate P budgets.

## Acknowledgments

We thank the captains and crews of the R/V Mediterranean Explorer and R/V Bat Galim, as well as the SoMMoS sampling team, for help with the sample collection. We also thank the captain, crew and science team of cruise HKB-SCS-2021, and especially Jiying Li, for their help in obtaining samples and data from the South China Sea. We thank Dikla Aharonovich for assistance throughout the project and Tom Reich for the biofilm images. This study was supported by grant 1786/20 from the Israel Science Foundation (to DS) and grant SMSEGL20SC02 from the Southern Marine Science and Engineering Guangdong Laboratory (Guangzhou, to DS). SG was supported by PhD scholarships from the Yochai Bin Nun foundation (IOLR) and the University of Haifa.

## Table and Figure legends

**Appendix S1**. Supporting Information.

**Table S1**. Macromolecular composition of heterotrophic bacteria and phytoplankton.

**Table S2**. Elemental composition of heterotrophic bacteria and phytoplankton.

**Table S3**. Comparison between P-RNA:P in different microorganisms.

**Fig. S1**. The effects of temperature on *Alteromonas* growth curve (A) and maximal growth rate (B) in 96 well plate batch cultures. All *Alteromonas* strains were grown in 96 well plates for 72h under temperature-controlled conditions: 22, 25, 26, 28, 30 and 32°C. Optical Density (OD at 600nm) was measured every 15 minutes, deep into the stationary phase. Each plot represents an average of three wells of a 96 well plate. Boxplots show the average and 75^th^ percentile, with outliers shown as circles. Blue branches on the schematic cladogram (based on phylogenetic distance) are *A. macleodii*, red branches are *A. mediterranea*.

**Fig. S2**. The effects of temperature on *Alteromonas* growth curves (A) and growth rate (B). Optical density (600nm) was measured every hour for 11 hours. The green data-points show when samples were collected for macromolecular analyses. Boxplots show the average and 75^th^ percentile.

**Fig. S3**. The effects of temperature on *Prochlorococcus* growth curves (A) and growth rate (B). Cells number were obtained from Flow Cytometry, and chlorophyll autofluorescence (Ex440/Em680) was measured every 2-3 days. The green time-points show when samples were collected for macromolecular analyses. Boxplots show the average and 75^th^ percentile.

**Fig. S4**. Biofilm formation by *Alteromonas*. (A) 5 *Alteromonas* strains after 24 hours and after 5 days. Experiment setup by Dikla Aharonovich and Tom Reich. Images taken by Tom Reich. (B) Comparison between OD absorbance and cells count of *Alteromonas* HOT1A3 and DE on the hours where samples were collected for macromolecular analysis (Fig S3). Note the difference in HOT1A3 between OD and cells count (∼3 fold).

**Fig. S5**. The effect of growth rate and cellular investment in RNA on RNA:protein ratio in *Alteromonas* and *Prochlorococcus*. (A) RNA per-cell quotas as a function of growth rate. (B) RNA:protein ratio as a function of growth rate. (C) RNA:protein ratio as a function of P-RNA:P. Note the different Y axis between *Alteromonas* and *Prochlorococcus*.

**Fig. S6**. Temperature effect on (A) P-RNA:P and (B) P-polyphosphate (P-poly:P). Concentration of polyphosphate for MIT9312 were below the limit of detection (<5uM) in this experiment. The different letters above the box plots in the graphs indicate statistically significant differences among the test temperatures (one-way ANOVA, *P*<0.05). NA= not available.

**Fig. S7**. Monthly changes in protein, RNA and POP concentrations, per-cell quotas, and ratios. (A) Protein; (B) RNA; (C) Particulate organic phosphorous (POP); (D) RNA:protein ratio; (E) Allocation of total cellular phosphorous to RNA (P-RNA:P). Blue; surface; Green; Deep chlorophyll maximum (DCM).

**Fig. S8**. Results from a gradient of anthropogenic input from Pearl River Estuary to the South China Sea (The HKB-SCS-2021 cruise, Jun-Jul 2021). (A) a map of the stations along the gradient. (B) Protein concentrations (N=1). (C) RNA concentrations (N=2). (D) Particulate organic phosphorous concentrations (N=1). (E) RNA:protein ratios (N=2). (F) Allocation of total cellular phosphorous to RNA (P-RNA:P).

**Fig. S9**. Difference in cell size between lab cultures and natural population of *Prochlorococcus* in the Eastern Mediterranean Sea. Cell size was estimated using flow cytometry (forward scatter, FSC). Samples from the EMS were taken from the deep chlorophyll maximum (DCM) (N=7 months in triplicates). Samples of *Prochlorococcus* cultures include both MIT9312 and NATL2A grown at 18°C (the temperature in the DCM) at different time points throughout the growth curves (N=72 in triplicates).

## Notes

### Competing Interest Statement

The authors have declared no competing interest.

## References

Acinas, S.G., Antón, J., and Rodríguez-Valera, F. (1999) Diversity of free-living and attached bacteria in offshore western Mediterranean waters as depicted by analysis of genes encoding 16S rRNA. Appl Environ Microbiol 65: 514–522.

Acker, M., Hogle, S.L., Berube, P.M., Hackl, T., Stepanauskas, R., Chisholm, S.W., and Repeta, D.J. (2022) Phosphonate production by marine microbes: exploring new sources and potential function. Proc Natl Acad Sci 119: e2113386119.

Aharonovich, D. and Sher, D. (2016) Transcriptional response of Prochlorococcus to co-culture with a marine Alteromonas: differences between strains and the involvement of putative infochemicals. ISME J 10: 2892–2906.

Ankrah, N.Y.D., May, A.L., Middleton, J.L., Jones, D.R., Hadden, M.K., Gooding, J.R., et al. (2014) Phage infection of an environmentally relevant marine bacterium alters host metabolism and lysate composition. ISME J 8: 1089–1100.

Basu, S., Bose, C., Ojha, N., Das, N., Das, J., Pal, M., and Khurana, S. (2015) Evolution of bacterial and fungal growth media. Bioinformation 11: 182–184.

Bertilsson, S., Berglund, O., Karl, D.M., and Chisholm, S.W. (2003) Elemental composition of marine Prochlorococcus and Synechococcus: Implications for the ecological stoichiometry of the sea. Limnol Oceanogr 48: 1721–1731.

Biller, S.J., Berube, P.M., Lindell, D., and Chisholm, S.W. (2015) Prochlorococcus: the structure and function of collective diversity. Nat Rev Microbiol 13: 13–27.

Biller, S.J., Coe, A., Roggensack, S.E., and Chisholm, S.W. (2018) Heterotroph Interactions Alter Prochlorococcus Transcriptome Dynamics during Extended Periods of Darkness. mSystems 3: 1–18.

Bru, S., Jimenez, J., Canadell, D., Arino, J., and Clotet, J. (2017) Improvement of biochemical methods of polyP quantification. Microb Cell 4: 6–15.

Casey, J.R., Boiteau, R.M., Engqvist, M.K.M., Finkel, Z. V, Li, G., Liefer, J., et al. (2022) Basin-scale biogeography of marine phytoplankton reflects cellular-scale optimization of metabolism and physiology. Sci Adv 8: eabl4930.

Cermak, N., Becker, J.W., Knudsen, S.M., Chisholm, S.W., Manalis, S.R., and Polz, M.F. (2017) Direct single-cell biomass estimates for marine bacteria via Archimedes’ principle. ISME J 11: 825–828.

Chomczynski, P. and Sacchi, N. (1987) Single-step method of RNA isolation by acid guanidinium thiocyanate-phenol-chloroform extraction. Anal Biochem 1: 581–585.

Christ, J.J. and Blank, L.M. (2018) Analytical polyphosphate extraction from Saccharomyces cerevisiae. Anal Biochem 563: 71–78.

Chrzanowski, T.H. and Grover, J.P. (2008) Element content of Pseudomonas fluorescens varies with growth rate and temperature: A replicated chemostat study addressing ecological stoichiometry. Limnol Oceanogr 53: 1242–1251.

Cotner, J.B., Makino, W., and Biddanda, B.A. (2006) Temperature affects stoichiometry and biochemical composition of Escherichia coli. Microb Ecol 52: 26–33.

Daines, S.J., Clark, J.R., and Lenton, T.M. (2014) Multiple environmental controls on phytoplankton growth strategies determine adaptive responses of the N:P ratio. Ecol Lett 17: 414–425.

Ben Ezra, T., Krom, M.D., Tsemel, A., Berman-Frank, I., Herut, B., Lehahn, Y., et al. (2021) Seasonal nutrient dynamics in the P depleted Eastern Mediterranean Sea. Deep Sea Res Part I Oceanogr Res Pap 176: 103607.

Farewell, A. and Neidhardt, F.C. (1998) Effect of temperature on in vivo protein synthetic capacity in Escherichia coli. J Bacteriol 180: 4704–4710.

Fraser, K.P.P., Clarke, A., and Peck, L.S. (2002) Low-temperature protein metabolism: seasonal changes in protein synthesis and RNA dynamics in the Antarctic limpet Nacella concinna Strebel 1908. J Exp Biol 205: 3077–3086.

Garcia-Martinez, J., Acinas, S.G., Massana, R., and Rodriguez-Valera, F. (2002) Prevalence and microdiversity of Alteromonas macleodii-like microorganisms in different oceanic regions. Environ Microbiol 4: 42–50.

Garcia, N.S., Bonachela, J.A., and Martiny, A.C. (2016) Interactions between growth-dependent changes in cell size, nutrient supply and cellular elemental stoichiometry of marine Synechococcus. ISME J 10: 2715–2724.

Geider, R.J. and La Roche, J. (2002) Redfield revisited: Variability of C:N:P in marine microalgae and its biochemical basis. Eur J Phycol 37: 1–17.

Giovannoni, S.J. (2017) SAR11 Bacteria: The Most Abundant Plankton in the Oceans. Ann Rev Mar Sci 9: 231–255.

Hanegraaf, P.P.F. and Muller, E.B. (2001) The Dynamics of the Macromolecular Composition of Biomass. J Theor Biol 212: 237–251.

Hui, S., Silverman, J.M., Chen, S.S., Erickson, D.W., Basan, M., Wang, J., et al. (2015) Quantitative proteomic analysis reveals a simple strategy of global resource allocation in bacteria. Mol Syst Biol 11: 784.

Hunter-cevera, K.R., Neubert, M.G., Solow, A.R., Olson, R.J., and Shalapyonok, A. (2014) Diel size distributions reveal seasonal growth dynamics of a coastal phytoplankter. Proc Natl Acad Sci 111: 1–6.

Ilikchyan, I.N., Mckay, R.M.L., Zehr, J.P., Dyhrman, S.T., and Bullerjahn, G.S. (2009) Detection and expression of the phosphonate transporter gene phnD in marine and freshwater picocyanobacteria. Environ Microbiol 11: 1314–1324.

Inomura, K., Omta, A.W., Talmy, D., Bragg, J., Deutsch, C., and Follows, M.J. (2020) A Mechanistic Model of Macromolecular Allocation, Elemental Stoichiometry, and Growth Rate in Phytoplankton. Front Microbiol 11: 1–22.

Jeffrey, W., Von Haven, R., Hoch, M., and Coffin, R. (1996) Bacterioplankton RNA, DNA, protein content and relationships to rates of thymidine and leucine incorporation. Aquat Microb Ecol 10: 87–95.

Kemp, P.F., Lee, S., and LaRoche, J. (1993) Estimating the growth rate of slowly growing marine bacteria from RNA content. Appl Environ Microbiol 59: 2594–2601.

Kirchman, D., K’nees, E., and Hodson, R. (1985) Leucine incorporation and its potential as a measure of protein synthesis by bacteria in natural aquatic systems. Appl Environ Microbiol 49: 599–607.

Kirchman, D.L. (2016) Growth Rates of Microbes in the Oceans. Ann Rev Mar Sci 8: 285–309.

Kulaev, I.S., Vagabov, V.M., and Kulakovskaya, T. V. (2005) The biochemistry of inorganic polyphosphates.

Lasternas, S., Agustí, S., and Duarte, C. (2010) Phyto- and bacterioplankton abundance and viability and their relationship with phosphorus across the Mediterranean Sea. Aquat Microb Ecol 60: 175–191.

Lee, S. and Fuhrman, J.E.D.A. (1987) Relationships between Biovolume and Biomass of Naturally Derived Marine Bacterioplankton. Appl Environ Microbiol 53: 1298–1303.

Lee, S.H. and Kemp, P.F. (1994) Single-cell RNA content of natural marine planktonic bacteria measured by hybridization with multiple 16S rRNA-targeted fluorescent probes. Limnol Oceanogr 39: 869–879.

Lin, Y., Gazsi, K., Lance, V.P., Larkin, A.A., Chandler, J.W., Zinser, E.R., and Johnson, Z.I. (2013) In situ activity of a dominant Prochlorococcus ecotype (eHL-II) from rRNA content and cell size. Environ Microbiol 15: 2736–2747.

Litchman, E., de Tezanos Pinto, P., Edwards, K.F., Klausmeier, C.A., Kremer, C.T., and Thomas, M.K. (2015) Global biogeochemical impacts of phytoplankton: A trait-based perspective. J Ecol 103: 1384–1396.

Lopez-Lopez A, Bartual SG, Stal L, Onyshchenko O, R.-V.F. (2005) Genetic analysis of housekeeping genes reveals a deep-sea ecotype of Alteromonas macleodii in the Mediterranean Sea. Environ Microbiol 7: 649–659.

López-Pérez, M., Gonzaga, A., Martin-Cuadrado, A.B., López-García, P., Rodriguez-Valera, F., and Kimes, N.E. (2013) Intra- and Intergenomic Variation of Ribosomal RNA Operons in Concurrent Alteromonas macleodii Strains. Microb Ecol 65: 720–730.

López-Pérez, M., Gonzaga, A., Martin-Cuadrado, A.B., Onyshchenko, O., Ghavidel, A., Ghai, R., and Rodriguez-Valera, F. (2012) Genomes of surface isolates of Alteromonas macleodii: The life of a widespread marine opportunistic copiotroph. Sci Rep 2: 1–11.

Ma, L., Calfee, B.C., Morris, J.J., Johnson, Z.I., and Zinser, E.R. (2018) Degradation of hydrogen peroxide at the ocean’s surface: the influence of the microbial community on the realized thermal niche of Prochlorococcus. ISME J 12: 473–484.

Makino, W., Cotner, J.B., Sterner, R.W., and Elser, J.J. (2003) Are bacteria more like plants or animals? Growth rate and resource dependence of bacterial C : N : P stoichiometry. Funct Ecol 17: 121–130.

Malmstrom, R.R., Coe, A., Kettler, G.C., Martiny, A.C., Frias-Lopez, J., Zinser, E.R., and Chisholm, S.W. (2010) Temporal dynamics of Prochlorococcus ecotypes in the Atlantic and Pacific oceans. ISME J 4: 1252–1264.

Martiny, A.C., Coleman, M.L., and Chisholm, S.W. (2006) Phosphate acquisition genes in Prochlorococcus ecotypes: evidence for genome-wide adaptation. Proc Natl Acad Sci 103: 12552–12557.

Martiny, A.C., Ma, L., Mouginot, C., Chandler, J.W., and Zinser, E.R. (2016) Interactions between Thermal Acclimation, Growth Rate, and Phylogeny Influence Prochlorococcus Elemental Stoichiometry. PLoS One 11: e0168291.

Martiny, A.C., Pham, C.T.A., Primeau, F.W., Vrugt, J.A., Moore, J.K., Levin, S.A., and Lomas, M.W. (2013) Strong latitudinal patterns in the elemental ratios of marine plankton and organic matter. Nat Geosci 6: 279–283.

Martiny, A.C., Talarmin, A., Mouginot, C., Lee, J.A., Huang, J.S., Gellene, A.G., and Caron, D.A. (2016) Biogeochemical interactions control a temporal succession in the elemental composition of marine communities. Limnol Oceanogr 61: 531–542.

Menzel, D.W. and Corwin, N. (1965) The measurement of total phosphorus in seawater based on the liberation of organically bound fractions by persulfate oxidation 1. Limnol Oceanogr 10: 280–282.

Moore, C.M., Mills, M.M., Arrigo, K.R., Berman-Frank, I., Bopp, L., Boyd, P.W., et al. (2013) Processes and patterns of oceanic nutrient limitation. Nat Geosci 6: 701–710.

Moore, L.R., Coe, A., Zinser, E.R., Saito, M.A., Sullivan, M.B., Lindell, D., et al. (2007) Culturing the marine cyanobacterium Prochlorococcus. Limnol Oceanogr Methods 5: 353–362.

Van Mooy, B.A.S., Rocap, G., Fredricks, H.F., Evans, C.T., and Devol, A.H. (2006) Sulfolipids dramatically decrease phosphorus demand by picocyanobacteria in oligotrophic marine environments. Proc Natl Acad Sci 103: 8607–8612.

Moreno, A.R. and Martiny, A.C. (2018) Ecological Stoichiometry of Ocean Plankton. Ann Rev Mar Sci 10: 43–69.

Morris, J.J., Kirkegaard, R., Szul, M.J., Johnson, Z.I., and Zinser, E.R. (2008) Facilitation of robust growth of Prochlorococcus colonies and dilute liquid cultures by “helper” heterotrophic bacteria. Appl Environ Microbiol 74: 4530–4534.

Mouginot, C., Zimmerman, A.E., Bonachela, J.A., Fredricks, H., Allison, S.D., Van Mooy, B.A.S., and Martiny, A.C. (2015) Resource allocation by the marine cyanobacterium S ynechococcus WH8102 in response to different nutrient supply ratios. Limnol Oceanogr 60: 1634–1641.

Partensky, F. and Garczarek, L. (2010) Prochlorococcus: Advantages and Limits of Minimalism. Ann Rev Mar Sci 2: 305–331.

Phillips, K.N., Godwin, C.M., and Cotner, J.B. (2017) The effects of nutrient imbalances and temperature on the biomass stoichiometry of freshwater bacteria. Front Microbiol 8: 1–11.

Quigg, A., Finkel, Z. V., Irwin, A.J., Rosenthal, Y., Ho, T.Y., Reinfelder, J.R., et al. (2003) The evolutionary inheritance of elemental stoichiometry in marine phytoplankton. Nature 425: 291–294.

Redfield, A.C. (1934) On the proportions of organic derivatives in sea water and their relation to the composition of plankton. James Johnstone Meml Vol 176–192.

Redfield, A.C. (1958) The biological control of chemical factors in the environment. Am Sci 46: 230A–221.

Rees, T.A. V. and Raven, J.A. (2021) The maximum growth rate hypothesis is correct for eukaryotic photosynthetic organisms, but not cyanobacteria. New Phytol 230: 601–611.

Reich, T., Ben-Ezra, T., Belkin, N., Tsemel, A., Aharonovich, D., Roth-Rosenberg, D., … & Sher, D. (2021) Seasonal dynamics of phytoplankton and bacterioplankton at the ultra-oligotrophic southeastern Mediterranean Sea. bioRxiv.

Reich, P.B. and Oleksyn, J. (2004) Global patterns of plant leaf N and P in relation to temperature and latitude. Proc Natl Acad Sci U S A 101: 11001–11006.

Rhee, G.Y. (1973) A continuous culture study of phosphate uptake, growth rate and polyphosphate in Scenedesmus sp. 1. J Phycol 9: 495–506.

Ribalet, F., Swalwell, J., Clayton, S., Jiménez, V., Sudek, S., Lin, Y., et al. (2015) Light-driven synchrony of Prochlorococcus growth and mortality in the subtropical Pacific gyre. Proc Natl Acad Sci 112: 7–11.

Rocap, G., Distel, D.L., Waterbury, J.B., and Chisholm, S.W. (2002) Resolution of Prochlorococcus and Synechococcus Ecotypes by Using 16S-23S Ribosomal DNA Internal Transcribed Spacer Sequences. Appl Environ Microbiol 68: 1180–1191.

Roth-rosenberg, D., Aharonovich, D., Omta, A., Follows, M.J., and Sher, D. (2020) Dynamic macromolecular composition and high exudation rates in Prochlorococcus. bioRxiv.

Roth Rosenberg, D., Haber, M., Goldford, J., Lalzar, M., Aharonovich, D., Al-Ashhab, A., … & Sher, D. (2021) Particle-associated and free-living bacterial communities in an oligotrophic sea are affected by different environmental factors. Environ Microbiol 23: 4295–4308.

Schirrmeister, B.E., Dalquen, D.A., Anisimova, M., and Bagheri, H.C. (2012) Gene copy number variation and its significance in cyanobacterial phylogeny. BMC Microbiol 12: 177.

Simon, M. and Azam, F. (1989) Protein content and protein synthesis rates of planktonic marine bacteria. Mar Ecol Prog Ser 51: 201–213.

Smith, P.K., Krohn, R.I., Hermanson, G.T., Mallia, A.K., Gartner, F.H., Provenzano, M.D., et al. (1985) Measurement of protein using bicinchoninic acid. Anal Biochem 150: 76–85.

Sterner, R.. and Elser, J.. (2002) Ecological stoichiometry: the biology of elements from molecules to the biosphere.

Tanaka, T., Thingstad, T.F., Christaki, U., Colombet, J., Cornet-Barthaux, V., Courties, C., et al. (2011) Lack of P-limitation of phytoplankton and heterotrophic prokaryotes in surface waters of three anticyclonic eddies in the stratified Mediterranean Sea. Biogeosciences 8: 525–538.

Toseland, A., Daines, S.J., Clark, J.R., Kirkham, A., Strauss, J., Uhlig, C., et al. (2013) The impact of temperature on marine phytoplankton resource allocation and metabolism. Nat Clim Chang 3: 979–984.

Vaulot, D., Marie, D., Olson, R.J., and Chisholm, S.W. (1995) Growth of Prochlorococcus, a photosynthetic prokaryote, in the equatorial Pacific Ocean. Science (80-).

Waldbauer, J.R., Rodrigue, S., Coleman, M.L., and Chisholm, S.W. (2012) Transcriptome and Proteome Dynamics of a Light-Dark Synchronized Bacterial Cell Cycle. PLoS One 7:.

Wei, T., Simko, V., Levy, M., Xie, Y., Jin, Y., & Zemla, J. (2017) Package ‘corrplot.’ Statistician 56: e24.

Woods, H.A., Makino, W., Cotner, J.B., Hobbie, S.E., Harrison, J.F., Acharya, K., and Elser, J.J. (2003) Temperature and the chemical composition of poikilothermic organisms. Funct Ecol 17: 237–245.

Yvon-Durocher, G., Dossena, M., Trimmer, M., Woodward, G., and Allen, A.P. (2015) Temperature and the biogeography of algal stoichiometry. Glob Ecol Biogeogr 24: 562–570.

Yvon-Durocher, G., Schaum, C.-E., and Trimmer, M. (2017) The temperature dependence of phytoplankton stoichiometry: investigating the roles of species sorting and local adaptation. Front Microbiol 8: 1–14.

Zimmerman, A.E., Allison, S.D., and Martiny, A.C. (2014) Phylogenetic constraints on elemental stoichiometry and resource allocation in heterotrophic marine bacteria. Environ Microbiol 16: 1398–1410.

Zinser, E.R., Johnson, Z.I., Coe, A., Karaca, E., Veneziano, D., and Chisholm, S.W. (2007) Influence of light and temperature on Prochlorococcus ecotype distributions in the Atlantic Ocean. Limnol Oceanogr 52: 2205–2220.

Zohary, T., Herut, B., Krom, M.D., Fauzi C. Mantoura, R., Pitta, P., Psarra, S., et al. (2005) P-limited bacteria but N and P co-limited phytoplankton in the Eastern Mediterranean—a microcosm experiment. Deep Sea Res Part II Top Stud Oceanogr 52: 3011–3023.

Zubkov, M. V (2014) Faster growth of the major prokaryotic versus eukaryotic CO2 fixers in the oligotrophic ocean. Nat Commun 5: 1–6.

