## Supporting Information for "Testing the Growth Rate and Temperature Compensation Hypotheses in Marine Bacterioplankton"

#### Supporting text

*Preliminary experiments to assess the growth dynamics of diverse Alteromonas strains across temperatures*

Five *Alteromonas* strains, belonging to two clades (*A. macleodii* and *A. mediterranea*) were grown at multiple temperatures in 96 well plates to assess the effect of temperature on growth rate and on the shape of the growth curve (Fig. S1A). Prior to the experiments, all strains were acclimated to the tested temperature for 48h and maintained in ProMM media (Morris et al., 2008). Experiments were started by inoculating 195µl ProMM media at the appropriate temperature with 5µl of a starter culture at stationary phase, followed by growth in an EnSpireTM 2300 Multilabel Reader (Perkin Elmer) under controlled temperature and measurements every 15 minutes (with prior shaking of 5 sec) for 72h. Growth rate was extracted from the exponential trend line equation (i.e. x coefficient) of the exponential growth phase.

In all *Alteromonas* strains tested, the growth rate increased with temperature from 22°C to 30-32°C (Fig. S1B, *P*-value<0.05, Pearson correlation). In addition, detailed analysis of the growth curves revealed several intriguing observations. Firstly, the shapes of the growth curves differed for different strains (Fig. S1A). For four out of the five *Alteromonas* strains the growth curve shapes were in agreement with their phylogeny, differing between the *A. macleodii* and *A. meditteranea* clades. *A. macleodii* ATCC and HOT1A3 showed a rapid initial growth, followed by a relatively abrupt cessation of growth after about 8 hours (Fig. S1A). This was followed by a second, slower growth phase and an extended stationary phase. The OD during the stationary phase generally increased with temperature. In contrast, the two *A. mediterranea* strains revealed a smoother transition from exponential growth to stationary phase, and the OD during stationary phase decreased with temperature. *A. macleodii* strain BS11 revealed a growth curve shape more like the *A. mediterranea* strains (Fig. S1A). Secondly, the growth rate of *A. macleodii* was higher than that of *A. mediterranea*, again with the exception of strain BS11 (Fig. S1B, *p*-value<0.05, t-test). *A. macleodii* and *A. mediterranea* differ not only phylogenetically but also in terms of their natural habitat. *A. macleodii* are more associated with the surface water while *A. mediterranea*, formerly known as “Deep Ecotypes” of *A. macleodii*, and are believed to grow mainly in deeper water (Ivars-Martinez et al., 2008; Lopez-Lopezet al., 2005). Interestingly, *A. macleodii* strain BS11, although more closely related to ATCC and HOT1A3, revealed a growth phenotype more reminiscent of the *A. mediterranea*. This strain is known to be different both physiologically (López-Pérez et al., 2012) and genetically (López-Pérez *et al.*, 2012, 2017) from other *A. macleodii* strains. The faster growth rate of *A. macleodii* ATCC and HOT1A3 and their rapid shifts between different growth "stages" may suggest they are more "copiotrophic" than the *A. mediterranea* strains, responding faster to changes in environmental conditions. The increased OD at stationary stage may suggest they are better adapted to the higher temperatures close to the surface. Alternatively, these differences in growth curves may reflect increased biofilm formation in *A. macleodii* (see below).

Based on these results, *A. macleodii* strain HOT1A3 and *A. mediterranea* strain DE were selected for further analysis of the macromolecular structure across four temperatures - 18°C, 22°C, 26°C and 30°C. To obtain sufficient biomass for biochemical analyses, larger volumes were required in these experiments (1L, in 1L bottles). In agreement with the experiments in 96 well plates, the growth rates in the scaled-up experiments increased with temperature (Fig. S2). The growth rates were slightly lower than in the 96 well plate format (~0.3-0.6 $h^{-1}$ in large volume compared with 0.5-0.8 $h^{-1}$ in 96 well plate for *A. macleodii* HOT1A3, ~0.27-0.45 $h^{-1}$ compared with 0.3-0.6 $h^{-1}$ for *A. mediterranea* DE), possibly due to the higher surface-volume ratio of the 96 well plates leading to increased oxygenation.

*Biofilm formation by Alteromonas*

In nature, *Alteromonas* strains are often associated with particles (Acinas et al., 1999; Garcia-Martinez et al., 2002; Roth Rosenberg et al., 2021). Additionally, the tendency of *A. macleodii* to form biofilm has been documented by Bae et al (2011), which showed *A. macleodii* to be a dominant biofilm-forming bacteria disrupting seawater reverse osmosis processes. Macroscopic observations of the five *Alteromonas* strains showed that they all produced floating biofilm particles, although the size of the particles and the time it took them to appear reproducibly differed between the strains (Fig. S4A). Specifically, the *A. macleodii* strains showed a higher tendency to form floating biofilms compared with *A. mediterranea*.

These biofilms may affect the estimates of the per-cell protein and RNA quotas in two ways. Firstly, the aggregation of cells in biofilms may result in an underestimation of the total number of cells in *A. macleodii* HOT1A3 counted by flow cytometry, due to incomplete separation of the cells (i.e., multiple cells being counted as a single event). This is supported by the observation that while the OD was generally similar between the two strains at the time of sampling, the cell counts were on average ~3 fold higher for *A. mediterranea* DE compared to *A. macleodii* HOT1A3 (Fig. S4B). Secondly, biofilms are comprised of extracellular macromolecules, including RNA and proteins (Grasland *et al.*, 2003). We still do not know the composition of the biofilm produced by different *Alteromonas* strains, and to what extent it contributes to the measured cell quotas of these macromolecules. We have attempted to minimize these biases by vigorously vortexing the samples prior to flow cytometry, but complete separation of the cells from the biofilm is currently not possible. Therefore, we focused subsequent analyses on the ratios between the different macromolecules as a way to assess cellular resource allocation, keeping in mind the caveat that some of these resources may be extracellular, and/or that the per-cell quotas may be overestimated, primarily for strain HOT1A3 (e.g. the per-cell protein quotas of *A. macleodii* HOT1A3, ~200-500fg/cell, are higher than published values for *A. macleodii* Alt1C, 70 fg/cell (Zimmerman et al., 2014)). Nonetheless, the estimated RNA:protein ratio and P-RNA:P in Alt1C (0.16 and 20, respectively) are in agreement with our ranges (0.06-1.25 and 13-87, respectively, Table 1). Further work is needed in order to characterize the biofilm-forming capacity of *Alteromonas* strains *in-vitro* and determine to what extent this process is reminiscent of their ability to colonize particles *in-situ*.

### Supplementary Tables

**Table S1.** Macromolecular composition of heterotrophic bacteria and phytoplankton.

| Macromolecule | Elemental composition C:N:P | Organism | % in cell (dry weight) |
|---|---|---|---|
| **Carbohydrates** | 6: - : - [a] | *E. Coli* | 5[b] |
| | | Marine bacteria | 5.4[c d] |
| | | Phytoplankton and cyanobacteria | 15[c d], 5-45[a], 16-38[e] |
| **Proteins** | 4.4: 1.1: - [a] * | *E. Coli* | 55[f] |
| | | Marine bacteria | 48[d], 63[g] |
| | | Phytoplankton and cyanobacteria | 32[d], 30-65[a], 37-52[e] |
| **Nucleic acids** | 9.5: 3.7: 1[h] | *E. Coli* | 20[f] |
| | | Marine bacteria | 4.4 RNA[d], 0.3 DNA[d] |
| | | Phytoplankton and cyanobacteria | 8-11[e]<br>5.7 RNA[d], 1 DNA[d]<br>3-15 RNA[a], 0.5-3 DNA[a] |
| **Phospholipids** | 38: 0.4: 1[a] | *E. Coli* | 10[f] |
| | | Marine bacteria | 5.4[d] |
| | | Phytoplankton and cyanobacteria | 17.3[d], 10-50[a], 8-13[e], 15[i] |
| **Chlorophyll-*a*** | 55: 4: - [a] | Phytoplankton and cyanobacteria | 1[d], 0.2-5[a] |
| **ATP** | 10: 5: 3[a] | Phytoplankton and cyanobacteria | <0.1[a] |

*Green*; phytoplankton/cyanobacteria. *Blue*; heterotrophic bacteria. *Orange*; *E. Coli*. *Black*; general feature.

*We are ignoring protein phosphorylation.

[a](Geider and La Roche, 2002); Algae and cyanobacteria under both nutrient-replete and nutrient-limited conditions.

[b](Milo et al., 2010); *E. coli*.

[c](Cotner *et al.*, 2006); *E. Coli*.

[d](Finkel *et al.*, 2016); Microalgae (exponentially growing). Under stationary growth: the average dry weight of protein declined to 27.0%, carbohydrate increased to 21.8% and lipid to 22.5%. There was insufficient RNA and DNA data from the stationary phase of growth to draw conclusions (N=2).

[e](Vargas *et al.*, 1998); Nitrogen-fixing cyanobacteria under diazotrophic conditions. Exponential growth: highest levels of carbohydrate and nucleic acids. Upon entering the stationary growth phase: highest levels of proteins and lipids.

[f](Milo *et al.*, 2010); *E. coli*.

[g](Simon and Azam, 1989); Marine bacteria (field measurements).

[h](Sterner and Elser, 2002); Prokaryote.

[i](Daines *et al.*, 2014); Diverse phytoplankton cultures.

**Table S2.** Elemental composition of heterotrophic bacteria and phytoplankton.

| Element | % in cell | Main molecules | Allocation (%) of total cellular element (C/N/P) * |
|---|---|---|---|
| | | Carbohydrates | 17,34[b], 41-47[c], 20-70[d] |
| **C** | 30-60[a] | Lipids | 12-25[b], 27-45[c], 20-40[d] |
| | | Protein | 28-59[b], ~10-55[c], 28[e] |
| | | Proteins, | 53-105[b], 45-85[c], 37[e] |
| **N** | 7-14[a] | Nucleic acids | 5-12[b], 1.7-18[c], 7[e] |
| | | Photosynthetic components (chl-*a*) | 0.4-1.4[b], 2.4[c] |
| | | Nucleic acids | 24-69[b], 30[f], 11-87[e], 50-80[g] |
| **P** | 2[a] | Phospholipids | 1-3.5[b], <5[f], 10-15[g] |
| | | Polyphosphate | 14-46[b], 40[f] |

*Green*; phytoplankton/cyanobacteria. *Blue*; heterotrophic bacteria. *Orange*; *E. Coli*. *Black*; general feature.

** i.e. how much N is allocated to protein?*

[a](Fagerbakke et al.,1996); Marine bacteria.

[b](Casey *et al.*, 2022); N and P starvation, *Prochlorococcus.*

[c](Geider and La Roche, 2002); Algae and cyanobacteria under both nutrient-replete and nutrient-limited conditions.

[d](Liefer *et al.*, 2019); Diatoms.

[e](Zimmerman et al.,2014); Gammaproteobacteria.

[f](Rhee, 1973); P-replete *Scenedesmus* (green algae).

[g](Cotner *et al.*, 2006); *E. Coli.*

**Table S3.** Comparison between P-RNA:P in different microorganisms.

| Domain/phylum | Organism | P in RNA [%] | Temperature [˚C] | Reference |
|---|---|---|---|---|
| Cyanobacteria | *Synechococcus* sp. WH8102 | ~5-15 | 24 | (Mouginot *et al.*, 2015) |
| Cyanobacteria | *Synechococcus* sp. WH8102 | ~10-15 | 24 | (Garcia *et al.*, 2016) |
| Cyanobacteria | *Prochlorococcus* MED4 | ~20-30 | 21 | (Casey *et al.*, 2022) |
| Bacteria | *Argobacterium* | ~2-30 | 10, 15, 20, 25, and 30 | (Phillips *et al.*, 2017) |
| Bacteria | *Arthrobacter* | ~2-30 | 10, 15, 20, 25, and 30 | (Phillips *et al.*, 2017) |
| Bacteria | *Flavobacterium* | ~10-30 | 10, 15, 20, 25, and 30 | (Phillips *et al.*, 2017) |
| Bacteria | Freshwater bacterial community | 25-93 | 25 | (Makino and Cotner, 2004) |
| Bacteria | *Ruegeria* DSS-3 | 99.3 | 20 | (Zimmerman *et al.*, 2014) |
| Bacteria | *Oceanicola* HTCC2516 | 19.8 | 20 | (Zimmerman *et al.*, 2014) |
| Bacteria | *Pelagibaca* HTCC2601 | 74 | 20 | (Zimmerman *et al.*, 2014) |
| Bacteria | *Alteromonas* 1C | 19.8 | 20 | (Zimmerman *et al.*, 2014) |
| Bacteria | *Vibrio* 1A | 11 | 20 | (Zimmerman *et al.*, 2014) |
| Bacteria | *Vibrio* 2D | 9.1 | 20 | (Zimmerman *et al.*, 2014) |
| Bacteria | *Marinomonas* 241 | 16.3 | 20 | (Zimmerman *et al.*, 2014) |
| Bacteria | *Marinomonas* 340 | 103 | 20 | (Zimmerman *et al.*, 2014) |
| Bacteria | *Psychrobacter* 224 | 6.3 | 20 | (Zimmerman *et al.*, 2014) |
| Bacteria | *Psychrobacter* 119 | 1.3 | 20 | (Zimmerman *et al.*, 2014) |
| Bacteria | *Halomonas* 005 | 72.1 | 20 | (Zimmerman *et al.*, 2014) |
| Bacteria | *Halomonas* 146 | 7.7 | 20 | (Zimmerman *et al.*, 2014) |
| Bacteria | *Pseudomonas fluorescens* | ~30 | 14, 20, and 28 | (Chrzanowski and Grover, 2008) |
| Bacteria | *E. coli* | ~40-80 | 37 | (Woods *et al.*, 2003) |
| Bacteria | *E. coli* | ~40-70 | 21, 24, 28, 33, 38, and 42 | (Cotner *et al.*, 2006) |
| Eukaryote | *Parthenocissus tricuspidate* | 20 | 29 | (Robson et al., 1959) |
| Eukaryote | *Chlorella vulgaris* | 27 | 24 | (Nyholm, 1977) |
| Eukaryote | *Scenedesmus* | ~30 | 20 | (Rhee, 1973) |

### Supporting Figures

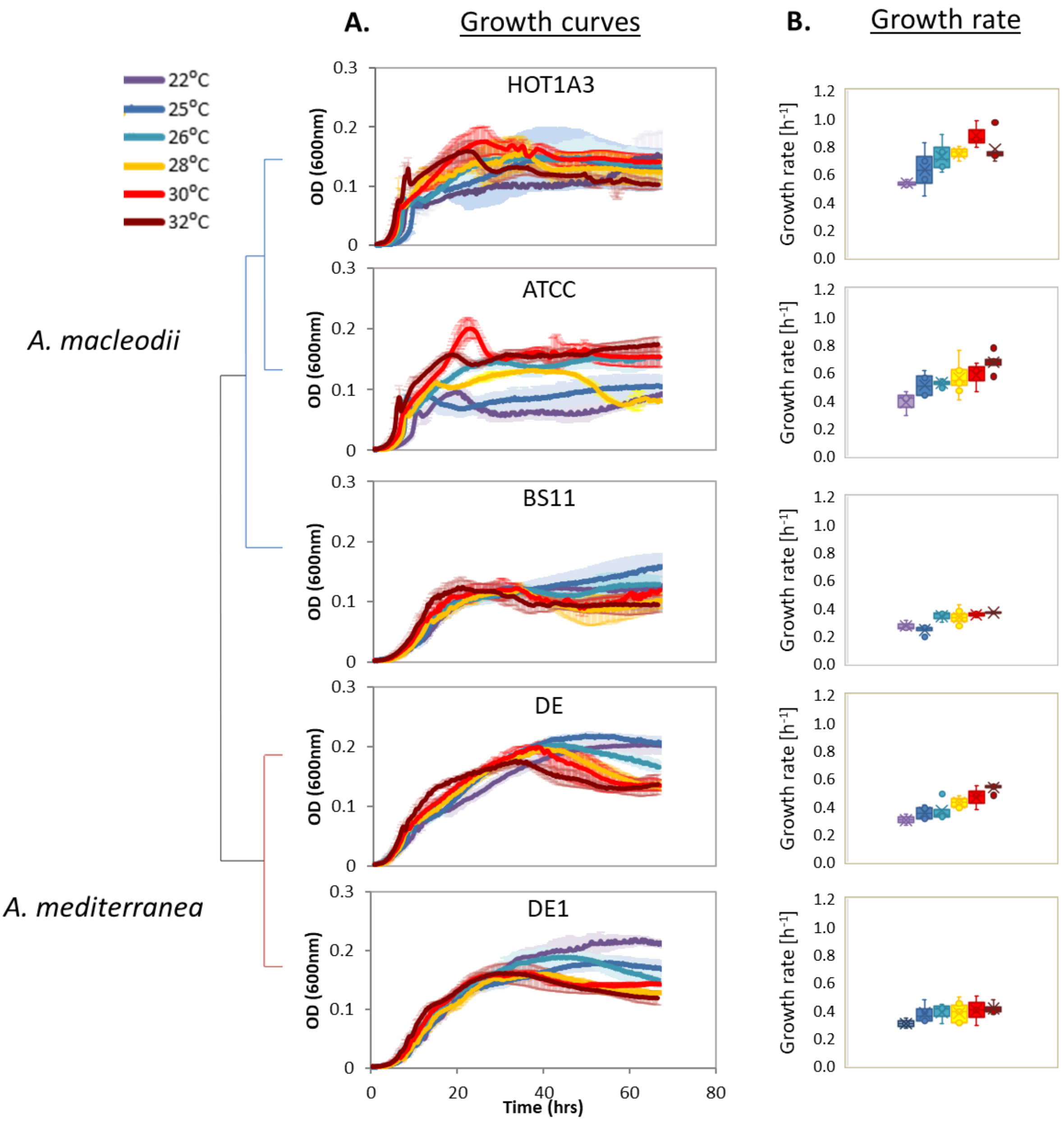


**Fig. S1**. The effects of temperature on *Alteromonas* growth curve (A) and maximal growth rate (B) in 96 well plate batch cultures. All *Alteromonas* strains were grown in 96 well plates for 72h under temperature-controlled conditions: 22, 25, 26, 28, 30 and 32°C. Optical Density (OD at 600nm) was measured every 15 minutes, deep into the stationary phase. Each plot represents an average of three wells of a 96 well plate. Boxplots show the average and 75$^{th}$ percentile, with outliers shown as circles. Blue branches on the schematic cladogram (based on phylogenetic distance) are *A. macleodii*, red branches are *A. mediterranea*.

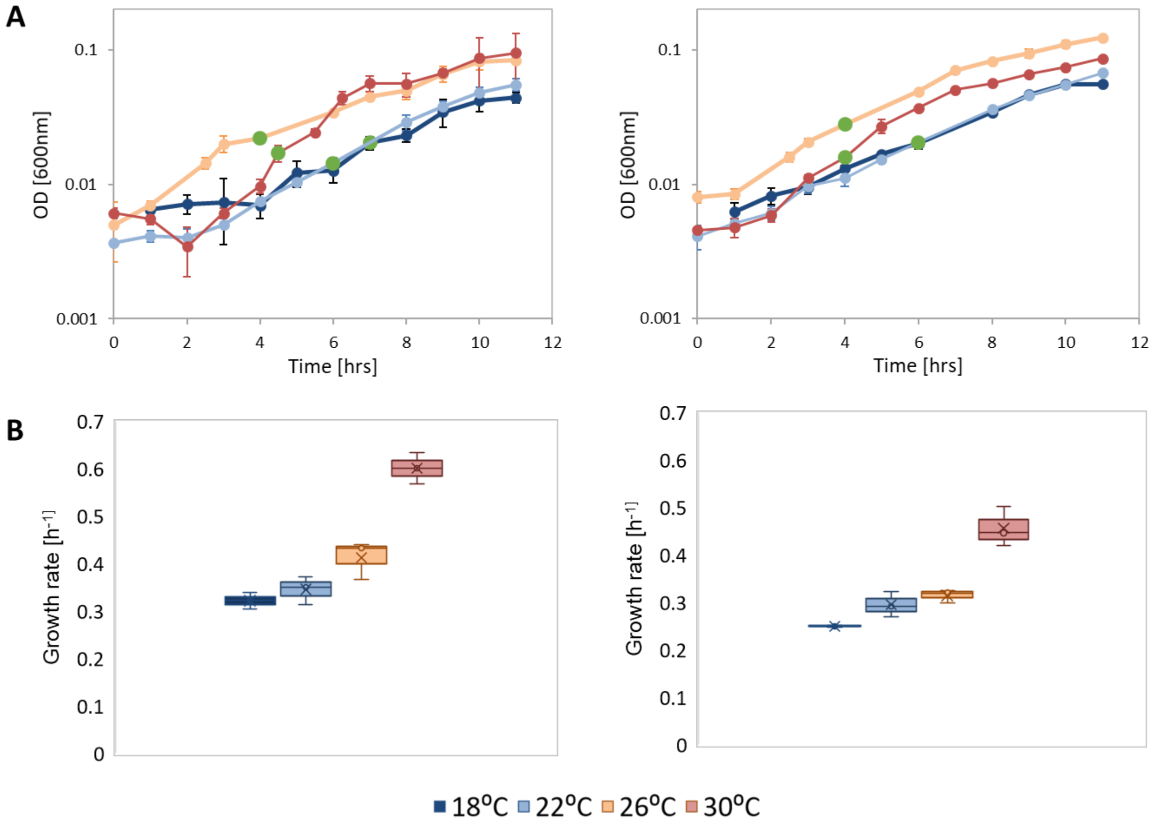


**Fig. S2**. The effects of temperature on *Alteromonas* growth curves (A) and growth rate (B). Optical density (600nm) was measured every hour for 11 hours. The green data-points show when samples were collected for macromolecular analyses. Boxplots show the average and 75$^{th}$ percentile.

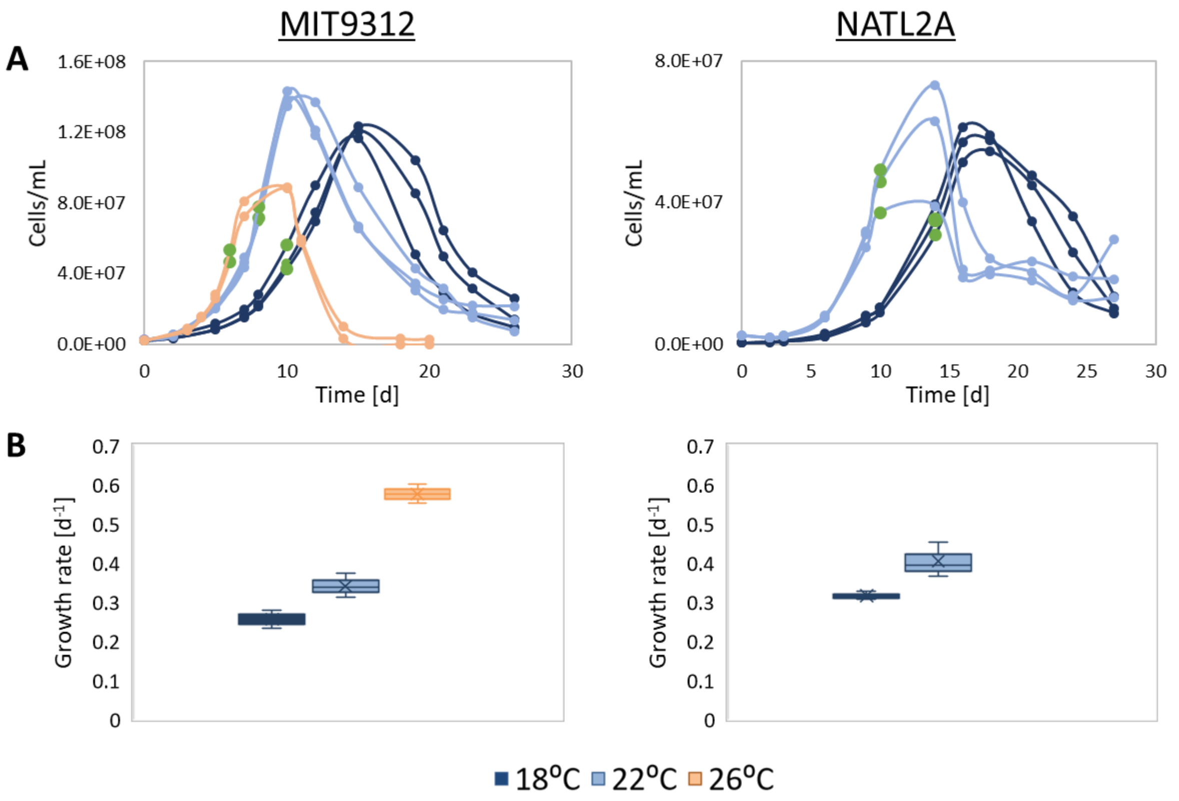


**Fig. S3**. The effects of temperature on *Prochlorococcus* growth curves (A) and growth rate (B). Cells number were obtained from Flow Cytometry, and chlorophyll autofluorescence (Ex440/Em680) was measured every 2-3 days. The green time-points show when samples were collected for macromolecular analyses. Boxplots show the average and 75$^{th}$ percentile.

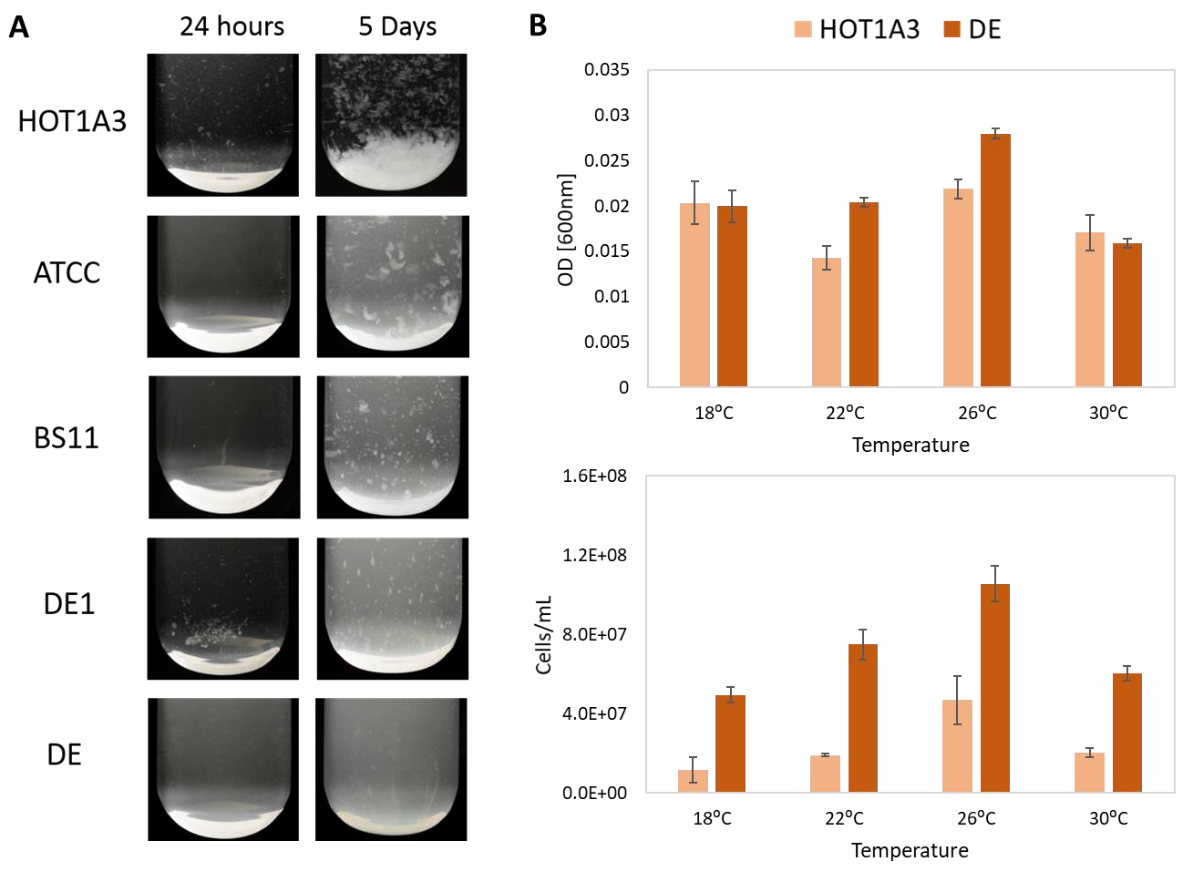


**Fig. S4.** Biofilm formation by *Alteromonas*. (A) 5 *Alteromonas* strains after 24 hours and after 5 days. Experiment setup by Dikla Aharonovich and Tom Reich. Images taken by Tom Reich. (B) Comparison between OD absorbance and cells count of *Alteromonas* HOT1A3 and DE on the hours where samples were collected for macromolecular analysis (Fig S3). Note the difference in HOT1A3 between OD and cells count (~3 fold).

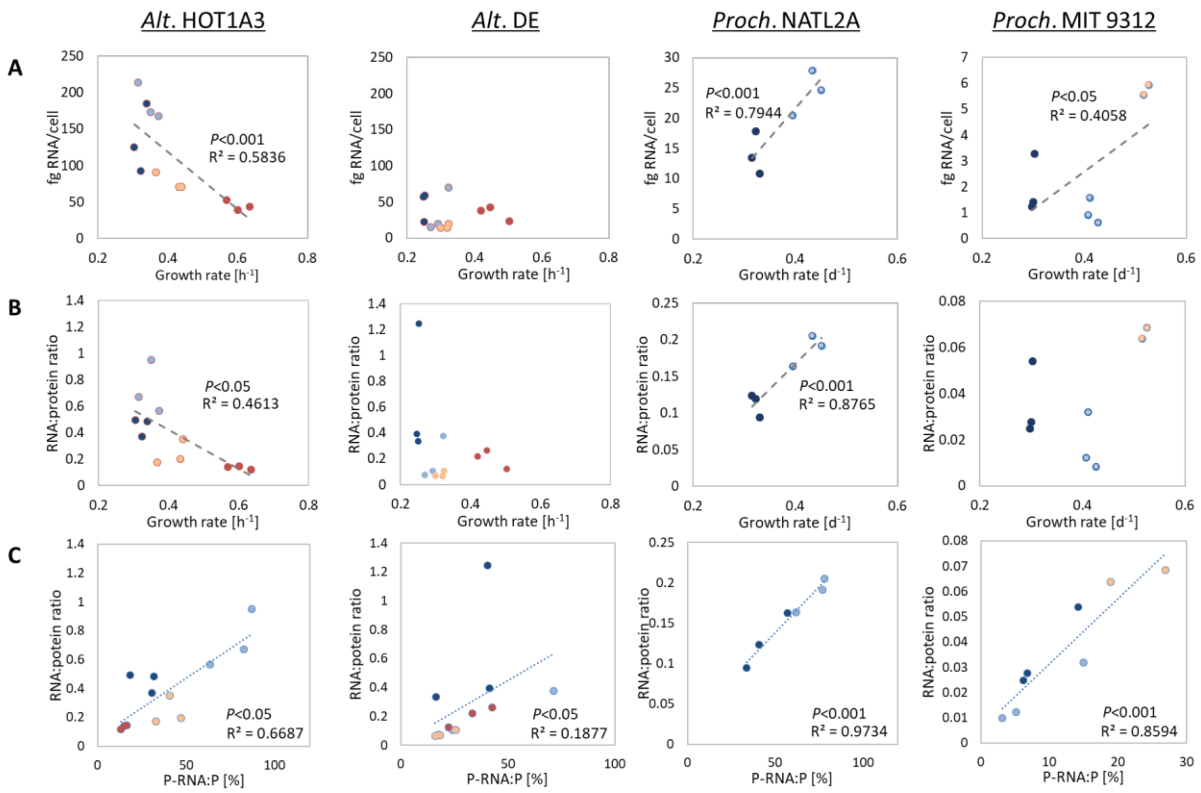


**Fig. S5**. The effect of growth rate and cellular investment in RNA on RNA:protein ratio in *Alteromonas* and *Prochlorococcus*. (A) RNA per-cell quotas as a function of growth rate. (B) RNA:protein ratio as a function of growth rate. (C) RNA:protein ratio as a function of P-RNA:P. Note the different Y axis between *Alteromonas* and *Prochlorococcus*.

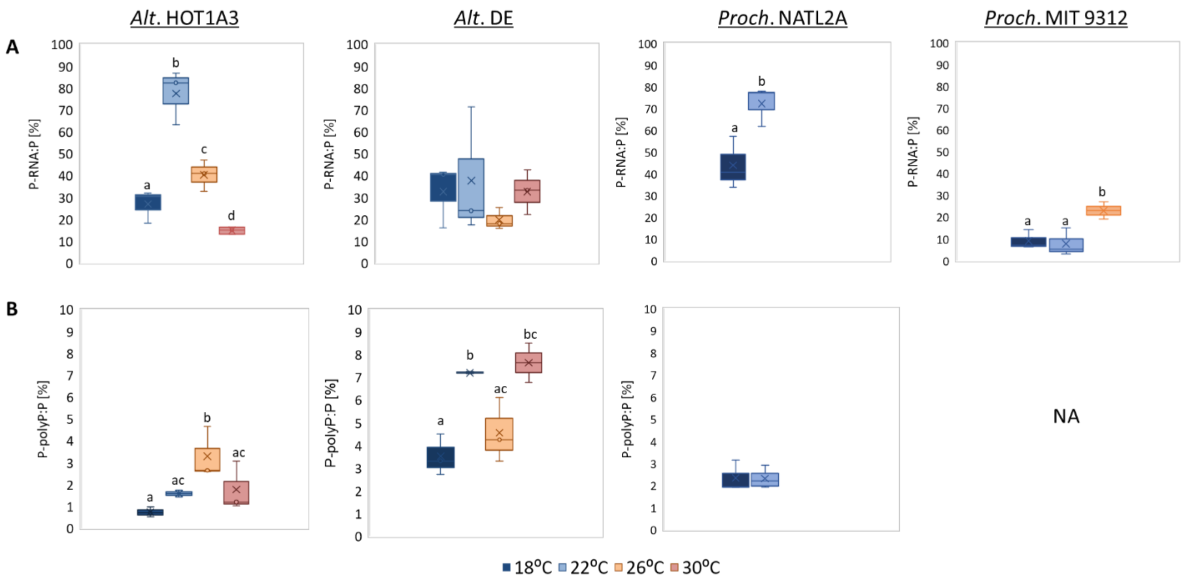


**Fig. S6**. Temperature effect on (A) P-RNA:P and (B) P-polyphosphate (P-poly:P). Concentration of polyphosphate for MIT9312 were below the limit of detection (<5uM) in this experiment. The different letters above the box plots in the graphs indicate statistically significant differences among the test temperatures (one-way ANOVA, *P*<0.05). NA= not available.

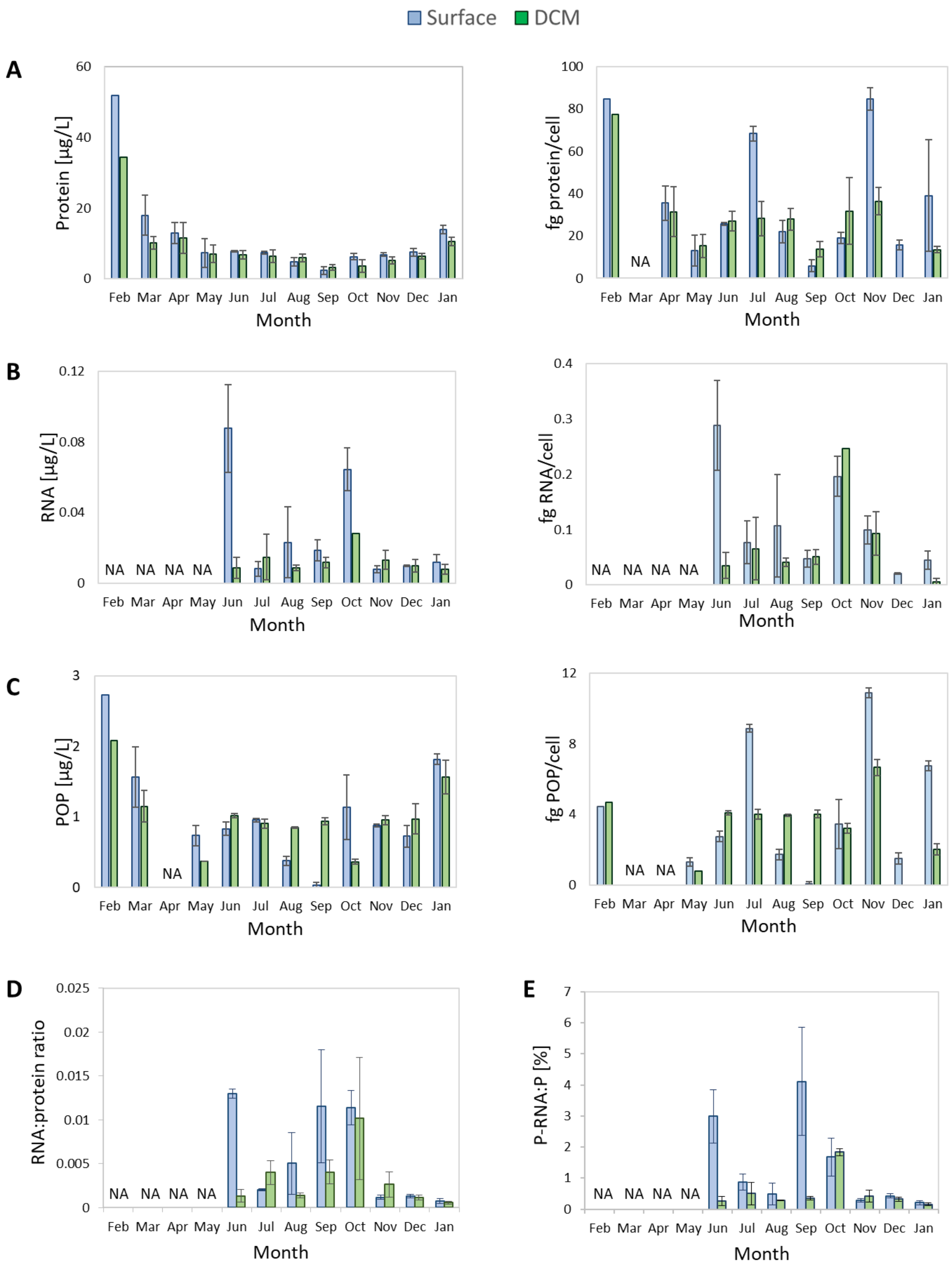


**Fig. S7**. Monthly changes in protein, RNA and POP concentrations, per-cell quotas, and ratios. (A) Protein; (B) RNA; (C) Particulate organic phosphorous (POP); (D) RNA:protein ratio; (E) Allocation of total cellular phosphorous to RNA (P-RNA:P). Blue; surface; Green; Deep chlorophyll maximum (DCM).

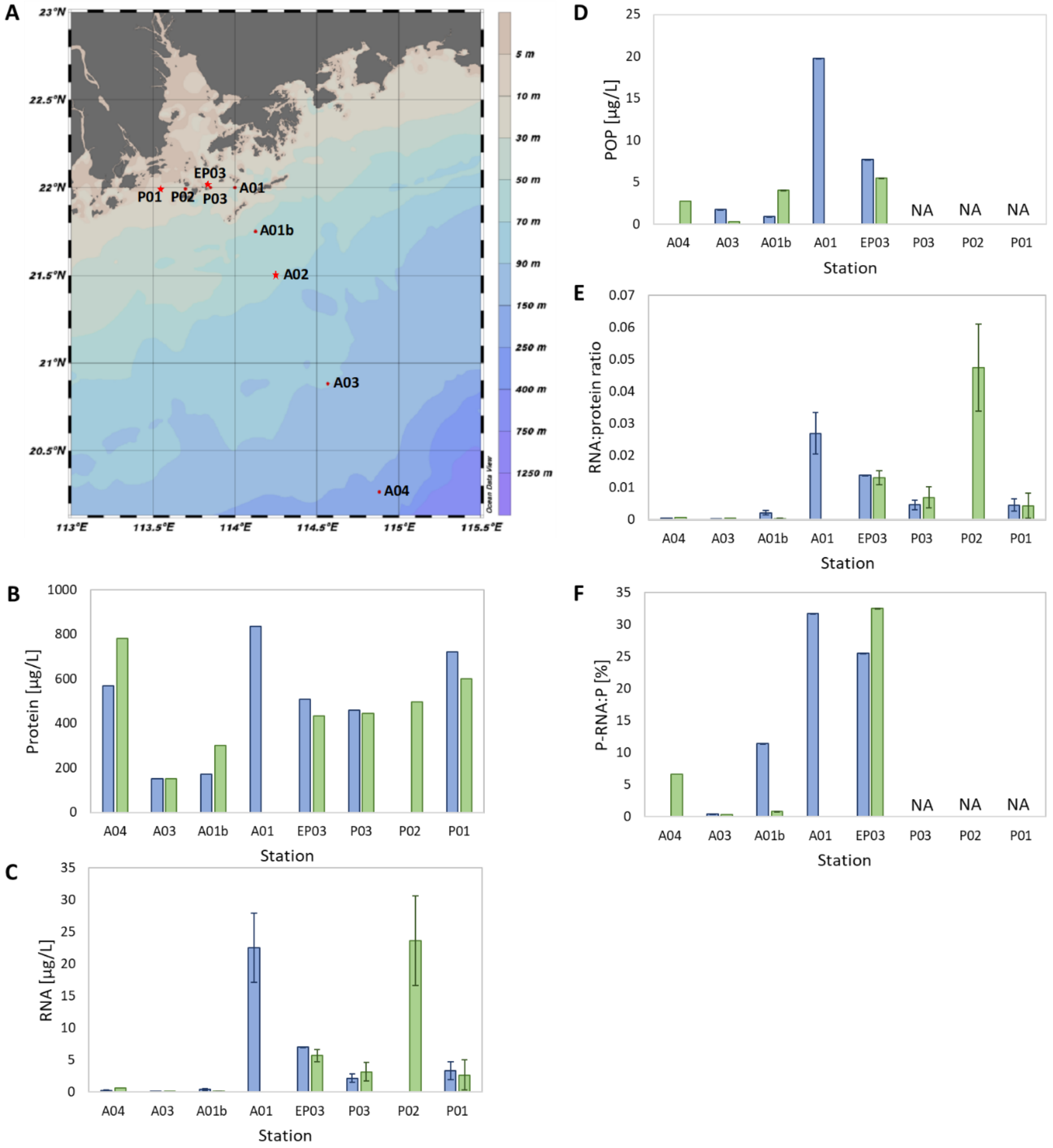


**Fig. S8.** Results from a gradient of anthropogenic input from Pearl River Estuary to the South China Sea (The HKB-SCS-2021 cruise, Jun-Jul 2021). (A) a map of the stations along the gradient. (B) Protein concentrations (N=1). (C) RNA concentrations (N=2). (D) Particulate organic phosphorous concentrations (N=1). (E) RNA:protein ratios (N=2). (F) Allocation of total cellular phosphorous to RNA (P-RNA:P).

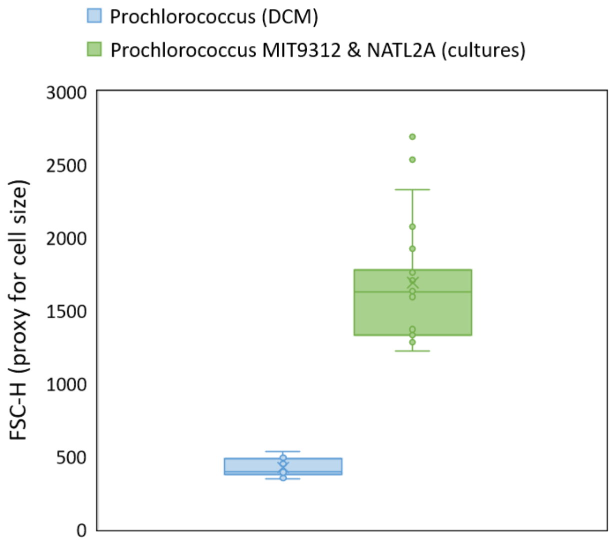


**Fig. S9.** Difference in cell size between lab cultures and natural population of *Prochlorococcus* in the Eastern Mediterranean Sea. Cell size was estimated using flow cytometry (forward scatter, FSC). Samples from the EMS were taken from the deep chlorophyll maximum (DCM) (N=7 months in triplicates). Samples of *Prochlorococcus* cultures include both MIT9312 and NATL2A grown at 18˚C (the temperature in the DCM) at different time points throughout the growth curves (N=72 in triplicates).

### References

Acinas, S.G., Antón, J., and Rodríguez-Valera, F. (1999) Diversity of free-living and attached bacteria in offshore western Mediterranean waters as depicted by analysis of genes encoding 16S rRNA. *Appl Environ Microbiol* **65**: 514–522.

Bae, H., Kim, H., Jeong, S., and Lee, S. (2011) Changes in the relative abundance of biofilm-forming bacteria by conventional sand-filtration and microfiltration as pretreatments for seawater reverse osmosis desalination. *Desalination* **273**: 258–266.

Casey, J.R., Boiteau, R.M., Engqvist, M.K.M., Finkel, Z. V, Li, G., Liefer, J., et al. (2022) Basin-scale biogeography of marine phytoplankton reflects cellular-scale optimization of metabolism and physiology. *Sci Adv* **8**: eabl4930.

Chrzanowski, T.H. and Grover, J.P. (2008) Element content of Pseudomonas fluorescens varies with growth rate and temperature: A replicated chemostat study addressing ecological stoichiometry. *Limnol Oceanogr* **53**: 1242–1251.

Cotner, J.B., Makino, W., and Biddanda, B.A. (2006) Temperature affects stoichiometry and biochemical composition of Escherichia coli. *Microb Ecol* **52**: 26–33.

Daines, S.J., Clark, J.R., and Lenton, T.M. (2014) Multiple environmental controls on phytoplankton growth strategies determine adaptive responses of the N:P ratio. *Ecol Lett* **17**: 414–425.

Fagerbakke, K.M., Heldal, M., and Norland, S. (1996) Content of carbon, nitrogen, oxygen, sulfur and phosphorus in native aquatic and cultured bacteria. *Aquat Microb Ecol* **10**: 15–27.

Finkel, Z. V., Follows, M.J., Liefer, J.D., Brown, C.M., Benner, I., and Irwin, A.J. (2016) Phylogenetic diversity in the macromolecular composition of microalgae. *PLoS One* **11**:.

Garcia-Martinez, J., Acinas, S.G., Massana, R., and Rodriguez-Valera, F. (2002) Prevalence and microdiversity of Alteromonas macleodii-like microorganisms in different oceanic regions. *Environ Microbiol* **4**: 42–50.

Garcia, N.S., Bonachela, J.A., and Martiny, A.C. (2016) Interactions between growth-dependent changes in cell size, nutrient supply and cellular elemental stoichiometry of marine Synechococcus. *ISME J* **10**: 2715–2724.

Geider, R.J. and La Roche, J. (2002) Redfield revisited: Variability of C:N:P in marine microalgae and its biochemical basis. *Eur J Phycol* **37**: 1–17.

Grasland, B., Mitalane, J., Briandet, R., Quemener, E., Meylheuc, T., Linossier, I., et al. (2003) Bacterial biofilm in seawater: cell surface properties of early-attached marine bacteria. *Biofouling* **19**: 307–313.

Ivars-Martinez, E., Martin-Cuadrado, A.-B., D'Auria, G., Mira, A., Ferriera, S., Johnson, J., et al. (2008) Comparative genomics of two ecotypes of the marine planktonic copiotroph Alteromonas macleodii suggests alternative lifestyles associated with different kinds of particulate organic matter. *ISME J* **2**: 1194–1212.

Liefer, J.D., Garg, A., Fyfe, M.H., Irwin, A.J., Benner, I., Brown, C.M., et al. (2019) The macromolecular basis of phytoplankton C:N:P under nitrogen starvation. *Front Microbiol* **10**: 1–16.

Lopez-Lopez A, Bartual SG, Stal L, Onyshchenko O, R.-V.F. (2005) Genetic analysis of housekeeping genes reveals a deep-sea ecotype of Alteromonas macleodii in the

Mediterranean Sea. *Environ Microbiol* **7**: 649–659.

López-Pérez, M., Gonzaga, A., Martin-Cuadrado, A.B., Onyshchenko, O., Ghavidel, A., Ghai, R., and Rodriguez-Valera, F. (2012) Genomes of surface isolates of Alteromonas macleodii: The life of a widespread marine opportunistic copiotroph. *Sci Rep* **2**: 1–11.

López-Pérez, M., Ramon-Marco, N., and Rodriguez-Valera, F. (2017) Networking in microbes: conjugative elements and plasmids in the genus Alteromonas. *BMC Genomics* **18**: 36.

Makino, W. and Cotner, J. (2004) Elemental stoichiometry of a heterotrophic bacterial community in a freshwater lake: implications for growth- and resource-dependent variations. *Aquat Microb Ecol* **34**: 33–41.

Milo, R., Jorgensen, P., Moran, U., Weber, G., and Springer, M. (2010) BioNumbers—the database of key numbers in molecular and cell biology. *Nucleic Acids Res* **38**: D750–D753.

Mouginot, C., Zimmerman, A.E., Bonachela, J.A., Fredricks, H., Allison, S.D., Van Mooy, B.A.S., and Martiny, A.C. (2015) Resource allocation by the marine cyanobacterium S ynechococcus WH8102 in response to different nutrient supply ratios. *Limnol Oceanogr* **60**: 1634–1641.

Nyholm, N. (1977) Kinetics of phosphate limited algal growth. *Biotechnol Bioeng* **19**: 467–492.

Phillips, K.N., Godwin, C.M., and Cotner, J.B. (2017) The effects of nutrient imbalances and temperature on the biomass stoichiometry of freshwater bacteria. *Front Microbiol* **8**: 1–11.

Rhee, G.Y. (1973) A continuous culture study of phosphate uptake, growth rate and polyphosphate in Scenedesmus sp. 1. *J Phycol* **9**: 495–506.

Robson, H. H., Budd, M. A., & Yost Jr, H.T. (1959) Comparison of the Nucleic Acid, Total Nitrogen, and Protein Nitrogen Levels of Normal and Tumor Tissue under Similar Growth Conditions. *Plant Physiol* **34**: 435–440.

Roth Rosenberg, D., Haber, M., Goldford, J., Lalzar, M., Aharonovich, D., Al-Ashhab, A., ... & Sher, D. (2021) Particle-associated and free-living bacterial communities in an oligotrophic sea are affected by different environmental factors. *Environ Microbiol* **23**:

4295–4308.

Simon, M. and Azam, F. (1989) Protein content and protein synthesis rates of planktonic marine bacteria. *Mar Ecol Prog Ser* **51**: 201–213.

Sterner, R.. and Elser, J.. (2002) Ecological stoichiometry: the biology of elements from molecules to the biosphere.

Vargas, M.A., Rodríguez, H., Moreno, J., Olivares, H., Del Campo, J.A., Rivas, J., and Guerrero, M.G. (1998) Biochemical composition and fatty acid content of filamentous nitrogen-fixing cyanobacteria. *J Phycol* **34**: 812–817.

Woods, H.A., Makino, W., Cotner, J.B., Hobbie, S.E., Harrison, J.F., Acharya, K., and Elser, J.J. (2003) Temperature and the chemical composition of poikilothermic organisms. *Funct Ecol* **17**: 237–245.

Zimmerman, A.E., Allison, S.D., and Martiny, A.C. (2014) Phylogenetic constraints on elemental stoichiometry and resource allocation in heterotrophic marine bacteria. *Environ Microbiol* **16**: 1398–1410.
